# The structure of the *Salmonella* phage epsilon15 tailspike reveals multiple O-antigen binding sites and a protruding esterase domain

**DOI:** 10.64898/2026.07.31.742079

**Authors:** Mateo Seoane-Blanco, Amaia Pereda, Nina Broeker, Michael McConnell, Francisco Javier Cañada, Stefanie Barbirz, Mark J. van Raaij

## Abstract

Many bacteriophages use tailspikes to degrade host bacterial polysaccharides, facilitating access to the outer membrane. The homotrimeric tailspikes of the *Salmonella* phage epsilon15 feature a slender phage-binding domain, a kink, and a barrel-shaped section with three petal-like protrusions. Here, we present the crystal structures of the monomeric protruding petal domain alone and of the trimeric barrel-shaped section with three petal domains. The barrel-shaped section includes a trimeric beta-helix, typical of phage tailspikes, alongside a trimeric beta-sandwich domain. The petal domain exhibits a fold characteristic of the serine-glycine-asparagine-histidine (SGNH) esterase family. Co-crystallisation with O-antigen fragments identified four binding sites on the tailspike: two adjacent sites on the surface of the triple beta-helix, one in the beta-sandwich domain and a fourth near the petal esterase site. These binding sites align with the expected orientation of the phage just before DNA transfer. Nuclear magnetic resonance spectroscopy and site-directed mutagenesis revealed an endorhamnosidase activity, showed that the reaction mechanism proceeds by inversion of the configuration and revealed that the active site is located at the junction of the two beta-helix binding sites. Analogous experiments also revealed an esterase site in the petal domain. Together, the structural and functional insights suggest a dual role for the phage epsilon15 tailspike: de-acetylation of the O-antigen, potentially affecting the local structure and lipopolysaccharide flexibility, plus cleavage of the O-antigen, enabling the phage to approach the bacterial membrane.

## 1. Introduction

Some bacteria hide their secondary phage receptors with thick layers of polysaccharides, such as lipopolysaccharide (LPS) or capsular polysaccharides. To overcome this barrier and approach the lipid membrane, adapted phages possess tailspikes that processively degrade them. In this way, tailspikes place the phage perpendicular to the cell envelope of the bacterium (Broeker & Barbirz, 2017; Figure 1*a*). These highly stable homotrimeric tailspikes (Mitraki *et al*., 2006) often have endoglycosidase activity (endorhamnosidase or endosialidase) or acetyl esterase activity and, by hydrolysing surface polysaccharides, they can have anti-biofilm activity (Prokhorov *et al*., 2017; Pires, Oliveira, *et al*., 2016).

**Figure 1.**
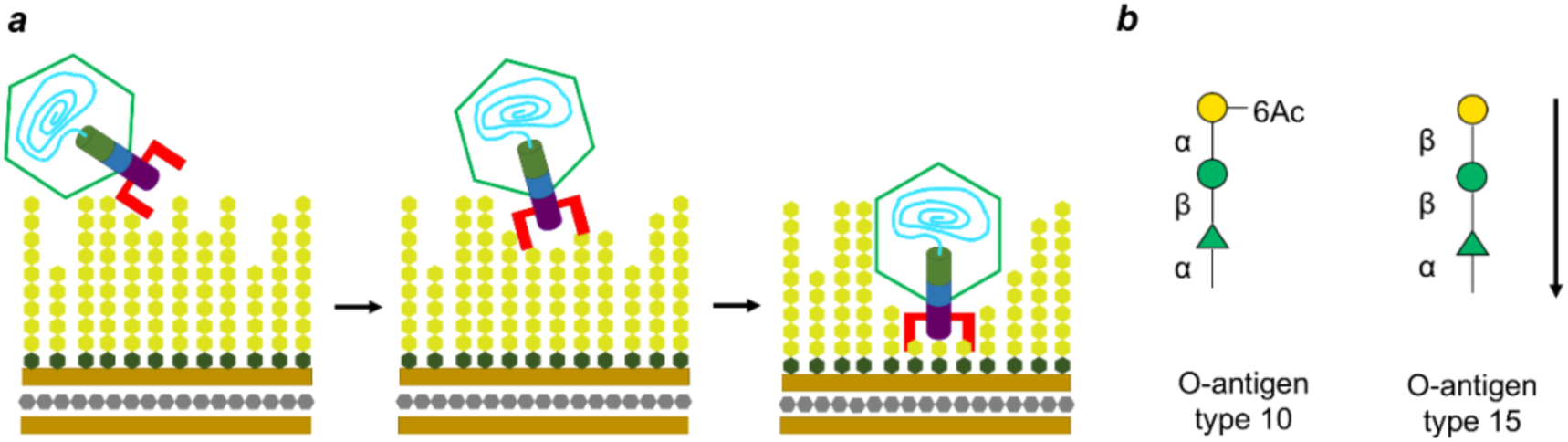
Infection of Salmonella by phage epsilone15 and nature of the O-antigen receptor. (*a*) Schematic model of a phage attachment and hydrolysis of its bacterial receptor. (*b*) Schematic representation of the trisaccharide repeating unit of the O-antigen type 10 present in the serovar Anatum and the O-antigen type present in the lysogenized serovar Anatum (the arrow indicates the non-reducing to reducing end direction). The Symbols Nomenclature For Glycans (Varki *et al*., 2015) was followed to represent the galactose (yellow circle) with an acetyl group attached to the carbon 6, the mannose (green circle) and the Rhamnose (green triangle).

Among phages that overcome bacterial polysaccharide barriers, *Salmonella* phage epsilon15 is a well-characterised example. This short-tailed bacteriophage (a podovirus) with an icosahedral head infects members of the Salmonella serogroup E1, such as *Salmonella enterica subsp. enterica* serovar Anatum A1 (Iseki & Sakai, 1953; Uetake *et al*., 1958; McConnell, et al., 1979; Sechter & Sechter-Mooreville, 1990; Jiang *et al*., 2006; Baker et al., 2013). Epsilon15 relies on six trimeric tailspikes, each made up of three copies of the gene product 20 (gp20), to bind and degrade the Anatum A1 O-antigen. These tailspikes, featuring a thin phage-binding domain, a kink, and a barrel-shaped domain resembling a three-petal flower (Jiang *et al*., 2006), play a central role in accessing the bacterial membrane, a process mediated by gp20’s enzymatic activity.

Epsilon15 tailspikes reversibly bind the Salmonella serogroup E1 (O:3,10) O-antigen type 10 (Figure 1*b*) (Kanegasaki & Wright, 1973; Sechter & Sechter-Mooreville, 1990), which consists of one to forty repeats of the partially acetylated trisaccharide –(D-Galp[6Ac/OH]-α-1–6-D-Manp-β-1–4-L-Rhap-α-1)– with this reducing-end Rha residue linked by α-1–3 bonds to the Gal residue of the next trisaccharide repeating unit (Robbins & Uchida, 1962; Gajdus *et al*., 2008; McConnell & Schoelz, 1983). Gp20 trimers specifically hydrolyse the alpha(1-3) link between the L-rhamnose and the D-galactose, generating hexasaccharides and larger oligosaccharides (Kanegasaki & Wright, 1973). This interaction is the crucial first step in the phage infection cycle, facilitating the attachment of the phage to the bacterial surface before genome injection. Understanding gp20’s enzymatic mechanisms is key to revealing how epsilon15 efficiently penetrates the bacterial defence and initiates infection.

In this study, we present the crystal structures of the epsilon15 tailspike gp20(248-1070), excluding the amino-terminal phage-binding domain, and the isolated petal domain, gp20(734-1070), both in apo forms and with bound O-antigen oligosaccharides. The petal domain exhibits esterase activity not previously described in this phage, deacetylating the galactose of LPS oligosaccharides. Gp20(248-1070) displays both endorhamnosidase and esterase activities, and furthermore, we find that the endorhamnosidase (assigned to Glycosyl hydrolase family 90, CAZY database, Drula *et al*., 2022) is acting through an inverting mechanism.

## 2. Materials and Methods

### 2.1 Cloning

Based on primary sequence and secondary structure prediction, seven gp20 constructs were designed to be cloned in pET28c(+). The first eight primers of the 0 were used to perform this task. A HindIII site exists between codons 200 and 287, necessitating the use of NotI for one of the reverse primers.

Then, three constructs without the petal domain were cloned into the plasmid pHTP1 system using a single ligase-independent reaction. The next 4 primers of 0 were used for this task.

The construct gp20(248-1070) was subcloned from the gp20(2-1070) construct based on sequence homology with the Det7 tailspike gp208ΔN, also called DettilonTSP due to its similarity with epsilon15 tailspike (Broeker *et al.,* 2019, PDB entry: 6F7D). The 13^th^ and 14^th^ primers of the 0 were used to carry out this task.

The gp20(248-778) construct was designed based on the structural knowledge from the petal domain and gp20ΔN, corresponding to the constructs gp20(734-1070) and gp20(248-1070). The gp20(248-778) construct was generated by deleting the C-terminal region of gp20(248-1070) in the pET28c(+) plasmid. The two last primers of 0 were used to perform this task.

The plasmid pET28c(+) encodes six-histidine tags (His-tags) just before and after the multi-cloning site, for inclusion at the N-and C-termini of the resultant expressed peptides. However, the sequence encoding the C-terminal His-tag was avoided by inserting a stop codon in front of it.

Construct gp20(434-1070) was ordered (GenScript) as a construct already inserted in the plasmid pET28a(+). Codons were optimised using the OPTIMIZER web server (Puigbò *et al*., 2007) with the codon list and frequencies for *Escherichia coli* B obtained from the website http://www.kazusa.or.jp/codon (Nakamura *et al*., 2000).

### 2.2. Mutagenesis

The mutants of gp20(248-1070) and gp20(734-1070) were generated using the NZYMutagenesis kit (NZYTech) according to supplier instructions and primers shown in the Table 2.

**Table 1.**
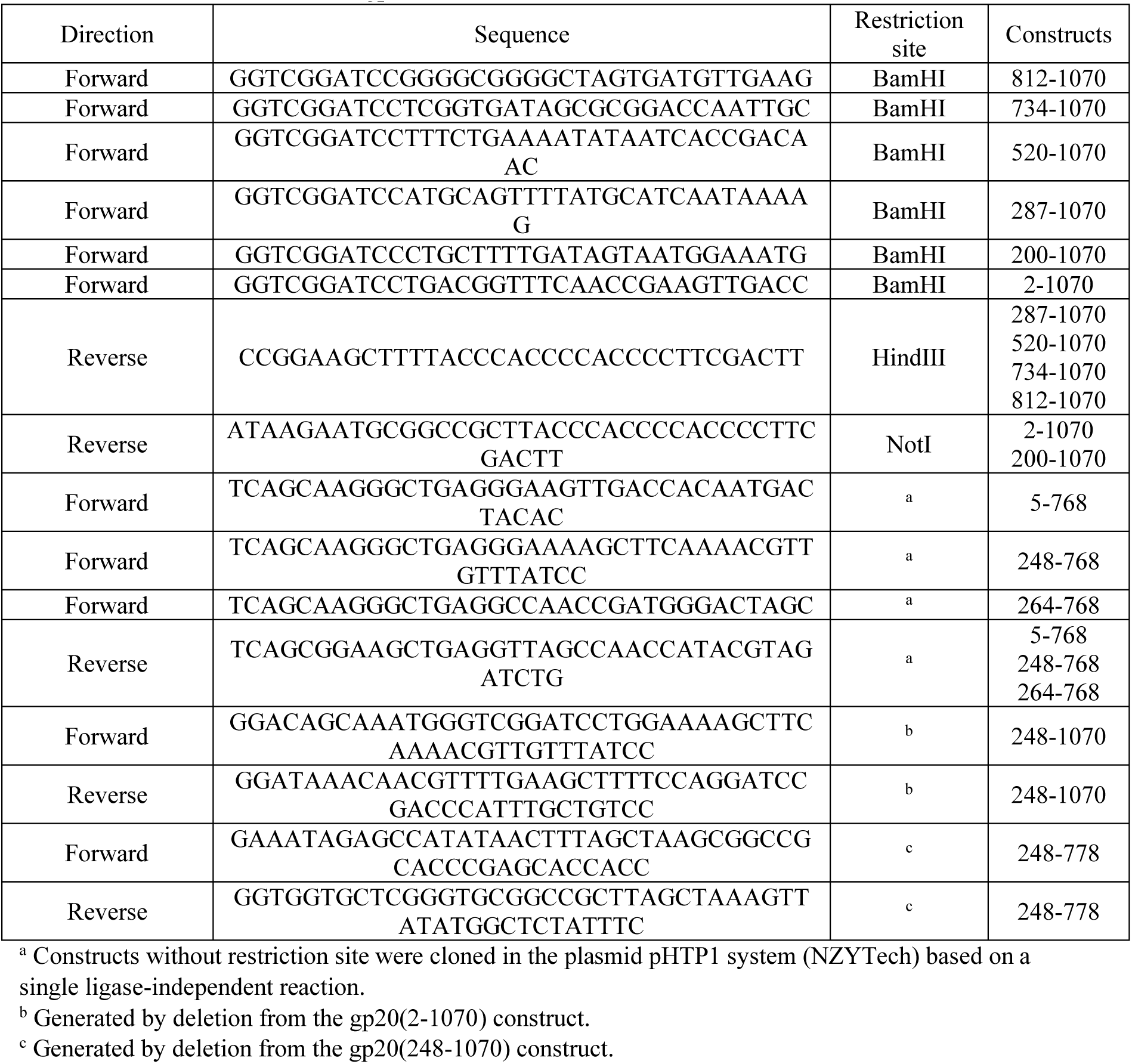
Primers used to clone gp20 constructs.

**Table 2.**
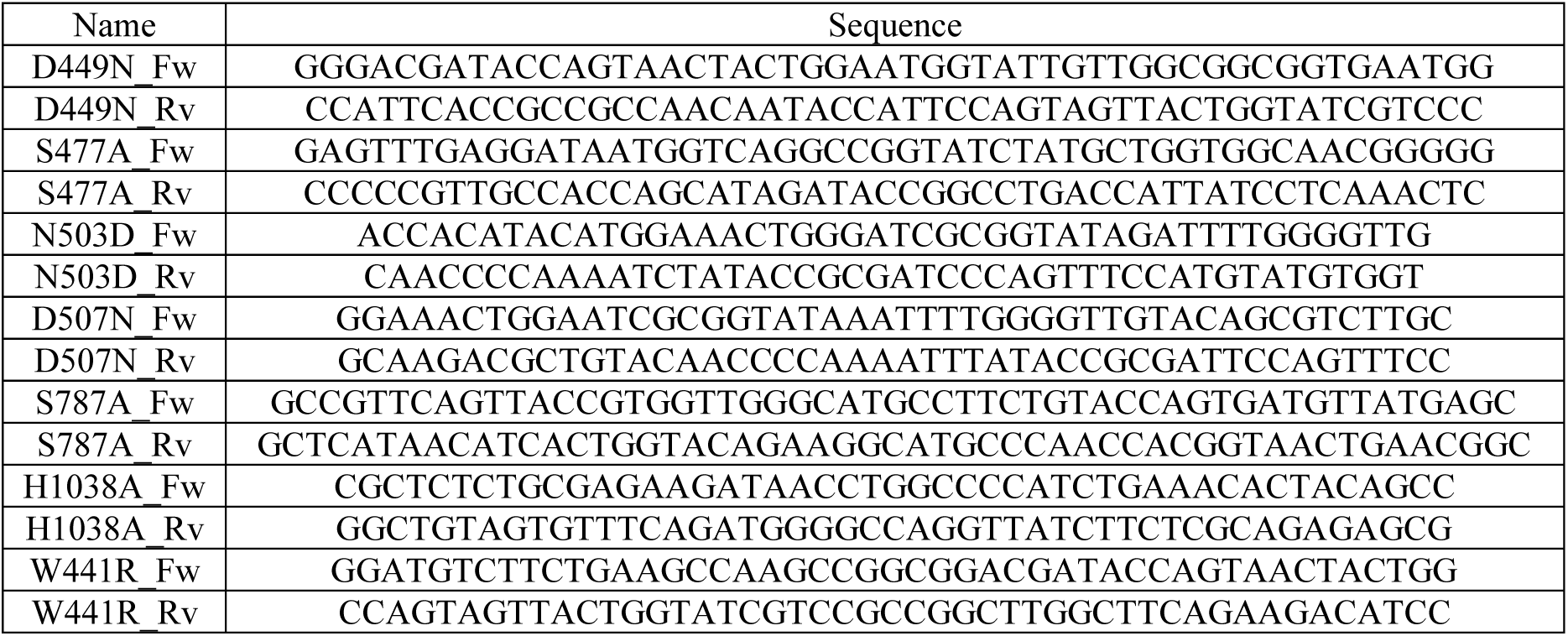
Primers used to generate the gp20(248-1070) and gp20(734-1070) mutants.

### 2.3. Expression and purification

For expression, constructs were transformed into the *Escherichia coli* strain BL21(DE3). When the culture reached an OD_600_=0.6-0.8, 1mM IPTG was added to induce protein expression at 25°C for gp20(248-1070) or at 15 °C for gp20(734-1070), gp20(248-778) and gp20(2-1070) overnight at 120 rpm. Cells were pelleted at 6000 x *g* for 10 minutes at 4 °C and resuspended in 20 mM Tris-HCl pH 7.5, 0.5 M NaCl, 5% (w/v) glycerol and 20 mM imidazole. Afterward, cells were sonicated and centrifuged at 15000 x *g* for 45 minutes at 4 °C. Later, the constructs were purified using immobilised metal ion affinity chromatography with nickel-nitrilotriacetic acid (Ni-NTA). Bound proteins were eluted with increasing concentrations of imidazole (from 20 to 1000 mM). Fractions containing the target proteins were pooled, thendialysed in cellulose membrane tubing against 20 mM Tris-HCl pH 7.5 and 0.2 M NaCl for 4 h at 4 °C and then, against 20 mM Tris-HCl pH7.5 overnight at 4 °C. The dialysed sample was centrifuged at 15000 x *g* for 10 minutes at 4 °C to remove aggregates. Then, ion exchange chromatography was performed using an ÄKTApurifier 10 FPLC system (Cytiva) with a RESOURCE^TM^ Q 6 mL column. Bound proteins were eluted with increasing concentrations of sodium chloride (0-1 M). Fractions with the pure proteins of interest were pooled, desalted and concentrated with centrifugal filters to 8 and 16 mg/mL for gp20(248-1070) and gp20(734-1070) respectively. Gp20(2-1070) and gp20(248-778) were concentrated to 1 mg/mL. Concentrated samples were centrifuged at 15000 x *g* for 10 minutes to remove aggregates.

Polysaccharide and oligosaccharide preparation from *S. anatum* LPS was as described previously (Zaccheus *et al*., 2012).

### 2.4. Crystallisation

Proteins were crystallised using the sitting-drop vapour-diffusion technique in MCR crystallisation plates (SWISSCI). Reservoirs were filled with 50 μL of crystallisation condition and lens with 1.5 μL (1 μL of protein and 0.5 μL of crystallisation condition). Due to ligand scarcity, only the protein drops with crystals were soaked with highly concentrated samples of Anatum O-antigen oligosaccharides. The petal domain crystals were obtained in 0.1 M Tris pH 8.0, 12% PEG 8000, selenomethionine-derivatised petal domain crystals in 0.1 M Citrate pH 5.5, 12% PEG 4000, and gp20(248-1070) in 3 M Na-formate. Drops of ligand were added to petal domain crystals in 0.1 M KCl, 0.1 M HEPES pH 7.0, 12% PEG 8000, and to gp20(248-1070) crystals in 2M Na-formate, 0.1 M Na-acetate pH 4.6. Petal domain crystals were soaked with ddH_2_O-dissolved Anatum O-antigen nonasaccharides at 20 mM for 17 h. Gp20(248-1070) crystals were soaked with ddH_2_O Anatum O-antigen hexasaccharides at 10 mM for 2 min.

### 2.5. Data collection

Crystals were diffracted at the XALOC beamline of the ALBA-CELLS synchrotron (Juanhuix *et al*., 2014). Diffraction image data sets were processed (indexed, integrated, merged and scaled) with AUTOPROC (Vonrhein *et al*., 2011) using the XALOC automated pipeline or inside the CCP4 suite (Winn *et al*., 2011) with iMOSFLM (Battyre *et al*., 2011) or XIA2/DIALS (Winn, 2010, Winn *et al*., 2018) and POINTLESS/AIMLESS (Evans, 2006, Evans & Murshudov, 2013). Datasets for selenomethionine derivatised gp20(734-1070) structures were phased with CRANK2 (Pannu *et al*., 2011). The native petal domain structure was obtained by molecular replacement with MOLREP (Vagin & Teplyakov, 2009) using the selenomethionine-derivatised model of the petal domain. In the gp20(248-1070) case, the selenomethionine-derivatised model of the petal domain together with the *Azotobacter vinelandii* Mannuronan C-5 epimerase AlgE6 A-module (PDB entry 5LW3; unpublished) were used for molecular replacement using PHASER (McCoy *et al*., 2007). Protein models were rebuilt with BUCCANEER (Cowtan, 2006) and COOT (Casañal *et al.,* 2020) and refined with REFMAC5 (Murshudov *et al*., 2011). The gp20(734-1070) and gp20(248-1070) models were obtained by molecular replacement using the apo versions of their respective constructs with MOLREP (Vagin & Teplyakov, 2009). Models were validated with MOLPROBITY (Williams *et al*., 2018). PyMOL (PYMOL Molecular Graphics System, Version 1.8 Schrödinger, LLC) and UCSF CHIMERA (Pettersen et al., 2004) were used for visualisation, analysis and preparation of protein model figures. The representations of the surface electrostatic potential were created with the PYMOL APBS plugin (Jurrus *et al*., 2018). Structure comparisons were performed using the DALI server (Holm, 2022), and oligomerisation parameters (accessible and buried surfaces, estimated dissociation energies) were analysed with PISA (Krissinel & Henrick, 2007).

### 2.6. Determination of carbohydrate-reducing ends

This assay measures the amount of sugar-reducing ends in a solution (Anthon & Barret, 2002, Zhang *et al*., 2011). The reagent 3-methyl-2-benzothiazolinone hydrazone hydrochloride (MBTH, Sigma Aldrich Scientific) reacts with reducing sugar ends (RSE), generating a chromogenic compound which has an absorption peak at 655 nm. We used MBTH to quantify O-antigen fragments produced by the E15 tailspike as it interacts with bacterial lipopolysaccharide (Anthon & Barret, 2002; Zhang *et al*., 2011). Briefly, 50 μL of tail protein (1 μM) and Anatum polysaccharide (1 mg/mL) were incubated for 2 h at 37 °C, before stopping the reaction with 50 μL of 0.25M NaOH. Then, 50 μL of a freshly prepared mixture of MBTH (1.5 mg/mL) and dithiothreitol(0.5mg/mL) was added to each tube (at neutral pH, the saccharide condenses with an MBTH molecule to form an adduct). The tubes were heated at 80 °C for 15 min, then treated with 100 μL of an oxidising solution containing 0.5% (w/v) FeNH4(SO4)2, 0.5% (w/v) sulfamic acid and 0.5M HCl. In acidic conditions, the adduct reacts with another MBTH molecule to form a highly coloured final product. Spectrophotometric measurements were performed in a Spectra Max ID3 reader (Molecular Devices), and a standard curve was generated using glucose at concentrations ranging from 25 to 1000 mM.

### 2.7. Electron microscopy of gp20(2-1070)

A 5-μL drop of gp20(2-1070) protein at 0.01 mg/mL was adsorbed for 5 minutes onto a glow-discharged carbon/collodion-coated copper grid (Gilder Grids), after which excess liquid was removed by quick blotting with filter paper (Whatman). The grid was washed twice with 2% (w/v) uranyl acetate, with blotting after each was. Finally, the grid was incubated with 2% (w/v) uranyl acetate for 2 min, blotted and air-dried on filter paper.

The grid was initially checked in a 100 kV JEOL JEM 1011 transmission electron microscope (JEOL) with a Gatan ES1000Ww camera (Gatan). Later, images were acquired with a FEI Tecnai FEG200 electron microscope (FEI) operated at 200 kV using a 4K x 4K Eagle CCD camera (FEI) at a 62000 x *g* magnification and a defocus range of 0.25-4 μm. The camera has a pixel size of 15 μm, giving a nominal sampling rate of pixel rate of 3.614 Å/pixel.

Images were processed using the Scipion Software Framework (de la Rosa-Trevín *et al*., 2016). The contrast transfer function was determined by CTFfind4 (Rohou & Grigorieff, 2015) for 441 images. Then, 29777 particles were manually picked with Xmipp3 – manual-picking software (Abrishami *et al*., 2013; de la Rosa-Trevín *et al*., 2013; Sorzano *et al*., 2013) in 110 x 110-pixel boxes. Bad particles were removed after being classified with the Xmipp3 – cl2d software (de la Rosa-Trevín *et al*., 2013; Sorzano *et al*., 2010; Sorzano *et al*., 2013). Then, the best averages were classified with the Relion – 2D classification software (Scheres *et al*., 2012; Zivanov et al., 2018) in one class and aligned with Xmipp3 – apply alignment 2d (de la Rosa-Trevín et al., 2013; Sorzano et al., 2013).

### 2.8. NMR spectroscopy

All experiments were recorded using 3 mm tubes (200 μL sample volume) and a Bruker AVIII-600 spectrometer equipped with a room temperature TXI probe head or a TXI cryoprobe, controlled by the software TOPSPIN (Bruker). 1D-1H and 2D TOCSY, HSQC spectra were recorded to assign the signals and confirm the structure. Diffusion ordered spectroscopy (DOSY) experiments (Groves *et al*., 2004) were performed to estimate the molecular size of the polysaccharide O-antigen samples and its fragments.

Bruker standard pulse program sequences included in TOPSIN were applied and spectra were processed using TOPSPIN 3.5 version. For ^1^H-NMR, pulse programs "zg", with 90° pulse, and "zgesgp", with water gradient suppression, were used. In case of TOCSY experiments, "dipsi2phpr" sequence was applied with mixing times (D9) of 20 and 70 ms and acquired with 12 ppm spectral window and 4096x256 data points. In the case of HSQC experiments, "hsqcedetgp" was applied with 10 ppm (1H) and 120 ppm (13C) spectral windows and acquired with 2048x256 data points. In the case of DOSY experiments, "ledbpgp2s" sequence was applied with 200 ms diffusion delay (D20) and 1.5 ms gradient length pulses (P30) and were acquired with 12 ppm spectral window and 16 gradient increments in the diffusion dimension.

Ligand-protein Sample preparation: The samples contained different variants of gp20 in which individual amino acids of the active site had been mutated or different domains had been removed. Depending on the experiment, samples were prepared between 0,5 to 2μM final concentration of protein in a buffer of 20mM potassium phosphate pH 7.5, 150mM NaCl in D2O and the polysaccharide was added to the required concentration (ligand:protein 50:1) from a 7 mg/mL (0,35mM, MW 20 kDa) stock solutions prepared in D2O.

On tube direct NMR monitoring of the enzymatic reaction progression: Different temperatures were tested: 278, 288 and 298 K, but spectra obtained at 298 K are presented. One dimensional experiments (^1^H-NMR, "zg" and "zgesgp") were recorded at 5-minute intervals for 24 hours, although in some exceptional cases, to obtain better results, experiments were performed every 2 minutes. The spectra were acquired with 32 scans and 32k points in the direct dimension. The time course experiments were also analysed with MNova software (Mestrelab Research, Spain).

## 3. Results

To obtain a soluble protein, we designed nine initial constructs, of which only two were found to be soluble. These constructs spanned amino acid ranges of 2-1070 and 734-1070 (Figure 3*h*). We note that this was done before the existence of artificial intelligence-based structure prediction or the knowledge of the DettilonTSP structure (Broeker *et al*., 2019); otherwise, we would most likely have had knowledge of the domain boundaries and more success in the design. Once the DettilonTSP structure became known, we also designed the gp20(248-1070) and gp20(248-778) constructs (Figure 3*h*), which were found to be soluble. First, the gp20(734-1070) construct, and later the gp20(248-1070) construct were successfully crystallised, allowing for structure determination. The non-soluble constructs had termini located within the petal domain (gp20(812-1070)), the beta-helix domain (520-1070, 434-1070, 287-1070, 264-768, and 200-1070), or the beta-sandwich domain (5-768, 248-768, and 264-768).

### 3.1 The epsilon15 tailspike structure reveals a novel petal domain

The longest crystallised fragment, gp20(248-1070), revealed three other domains: a beta-helix domain, a beta-sandwich domain, and a novel petal domain (Figure 3*a* & *e*). The beta-helix and the beta-sandwich domain are structurally homologous to the gp208 tailspike from phage Det7 (PDB entry: 6F7K, Broeker *et al.,* 2019) (Figure 3*b*). The beta-helix spans from residues 248 to 644 (279 to 667 in gp208) and the beta-sandwich covers residues 645 to 773 (668 to 798 in gp208). The primary difference between these homologous domains is the presence of a loop (456-459) in gp20, flanking the groove of the beta-helix domain. Notably, Glu456 binds to His382 on the opposite side of the groove (Figure 3*c*).

The trimer of beta-helices and beta-sandwiches forms the barrel-shaped core of the tailspike. In epsilon15, the final domain extends from this axis, creating three petal-like protrusions (Figure 3*d-e*). This petal domain spans residues 778 to 1070 and is connected to the beta-sandwich domain by a short linker. Unlike the other domains in the trimeric tailspike, the petal domains do not interact with each other, which led to the crystallisation of the shorter construct (residues 734-1070) as a monomer (Figure 3*f-g*). However, the residues corresponding to the beta-sandwich domain and the linker (734-777) were unresolved in the crystal structure. The petal domains in both crystal structures are almost structurally identical, with an RMSD of 0.4 Å.

Two constructs of gp20 were crystallised, one containing residues 734-1070, and the other containing residues 248-1070 (Figure 3*a* & *f)*. The longest fragment (residues 248-1070) diffracted to 1.94 Å (Table 3a), lacking the phage-binding domain. Attempts to crystallise the full-length tailspike were unsuccessful. Cryo-EM images revealed significant variability in the angle between the barrel-shaped body and the phage-binding domain. This angle ranged from 170° to 40°, with 60° being the most common, observed in 31% of the particles (Figure 2). This flexibility likely renders the full-length tailspike unsuitable for crystallisation.

**Table 3a.** Crystallographic statistics of gp20(734-1070) datasets

|  | gp20(734-1070)<br>SeMet | gp20(734-1070) | gp20(734-1070)<br>- ligand |
| --- | --- | --- | --- |
| **Data collection** |  |  |  |
| Radiation source | BL13-XALOC<br>(ALBA-CELLS) | BL13-XALOC<br>(ALBA-CELLS) | BL13-XALOC<br>(ALBA-CELLS) |
| Wavelength (Å) | 0.97852 | 0.97854 | 0.97980 |
| Detector | PILATUS 6M -<br>DECTRIS | PILATUS 6M -<br>DECTRIS | PILATUS 6M --<br>DECTRIS |
| Crystal-to-detector distance (mm) | 242.5 | 188.5 | 502.3 |
| **Data processing** |  |  |  |
| Space group | P2<sub>1</sub>2<sub>1</sub>2<sub>1</sub> | C2 | P2<sub>1</sub>2<sub>1</sub>2<sub>1</sub> |
| Cell edges (a, b, c; Å) | 41.7, 49.7, 144.7 | 154.2, 41.5, 49.4 | 41.5, 49.7, 144.4 |
| Cell angles (α, β, γ, °) | 90.0, 90.0, 90.0 | 90.0, 102.0, 90.0 | 90.0, 90.0, 90.0 |
| Resolution range (Å) | 144.69–1.64<br>(1.73-1.64) | 45.21-1.32<br>(1.40-1.32) | 144.42-2.09<br>(2.14-2.09) |
| Number of unique reflections | 37674 (5414) | 67546 (9663) | 17754 (2072) |
| Completeness (%) | 100.0 (100.0) | 95.0 (93.6) | 95.6 (78.8) |
| Anomalous completeness (%) | 100.0 (100.0) | - | - |
| Multiplicity | 12.8 (13.1) | 2.5 (2.5) | 10.4 (5.3) |
| Anomalous multiplicity | 6.8 (6.8) | - | - |
| CC 1/2<sup>a</sup> | 0.999 (0.781) | 0.999 (0.848) | 0.994 (0.599) |
| Resolution (Å) at CC<sub>anomfit</sub> = 0.15 | 1.96 | - | - |
| Wilson B factor (Å<sup>2</sup>) | 20.3 | 14.4 | 25.5 |
| <I/σ(I)> | 18.3 (2.4) | 13.2 (2.3) | 10.0 (1.9) |
| **Phasing** |  |  |  |
| Heavy atoms sites | 5 Se | - | - |
| Correlation coeff. (all/weak)<sup>b</sup> | 48.8/29.9 | - | - |
| Combined DM FOM + phasing<br>CLD score (chosen/rejected hand)<sup>c</sup> | 19.5/0.0 | - | - |
| **Refinement** |  |  |  |
| Resolution range (Å) | 72.35-1.64 | 45.21-1.32 | 72.21-2.09 |
| Reflections used | 37601 | 67546 | 17697 |
| Reflections used for R-free | 1882 | 3447 | 878 |
| R-factor/R-free<sup>d</sup> | 0.15/0.18 | 0.11/0.14 | 0.19/0.23 |
| **Model statistics** |  |  |  |
| Amino acid coverage | 778-1070 | 778-1070 | 778-1069 |
| Atoms<br>(protein/ions/ligands/water) | 2251/0/86/249 | 2270/0/ 72/366 | 2234/0/92/186 |
| Ramachandran<sup>e</sup><br>(%favoured/allowed/outliers) | 98.37/1.63/0.00 | 98.00/2.00/0.00 | 97.54/2.46/0.00 |
| RMSD (bonds, Å / angles, °)<sup>f</sup> | 0.0096/1.56 | 0.0052/1.38 | 0.0029/1.03 |
| B-factor<br>(protein/ions/<br>ligands/water) | 19.93/0/50.22/34.<br>52 | 14.58/-<br>/37.87/33.50 | 24.08/-<br>/45.04/30.12 |
| MolProbity score/percentile<sup>g</sup> | 1.18/98<sup>th</sup> | 1.23/95<sup>th</sup> | 1.05/100<sup>th</sup> |
| Clashscore | 1.94 | 2.58 | 1.33 |
| PDB entry | 9SIJ | 9SIK | 9SIL |

**Table 3b.**
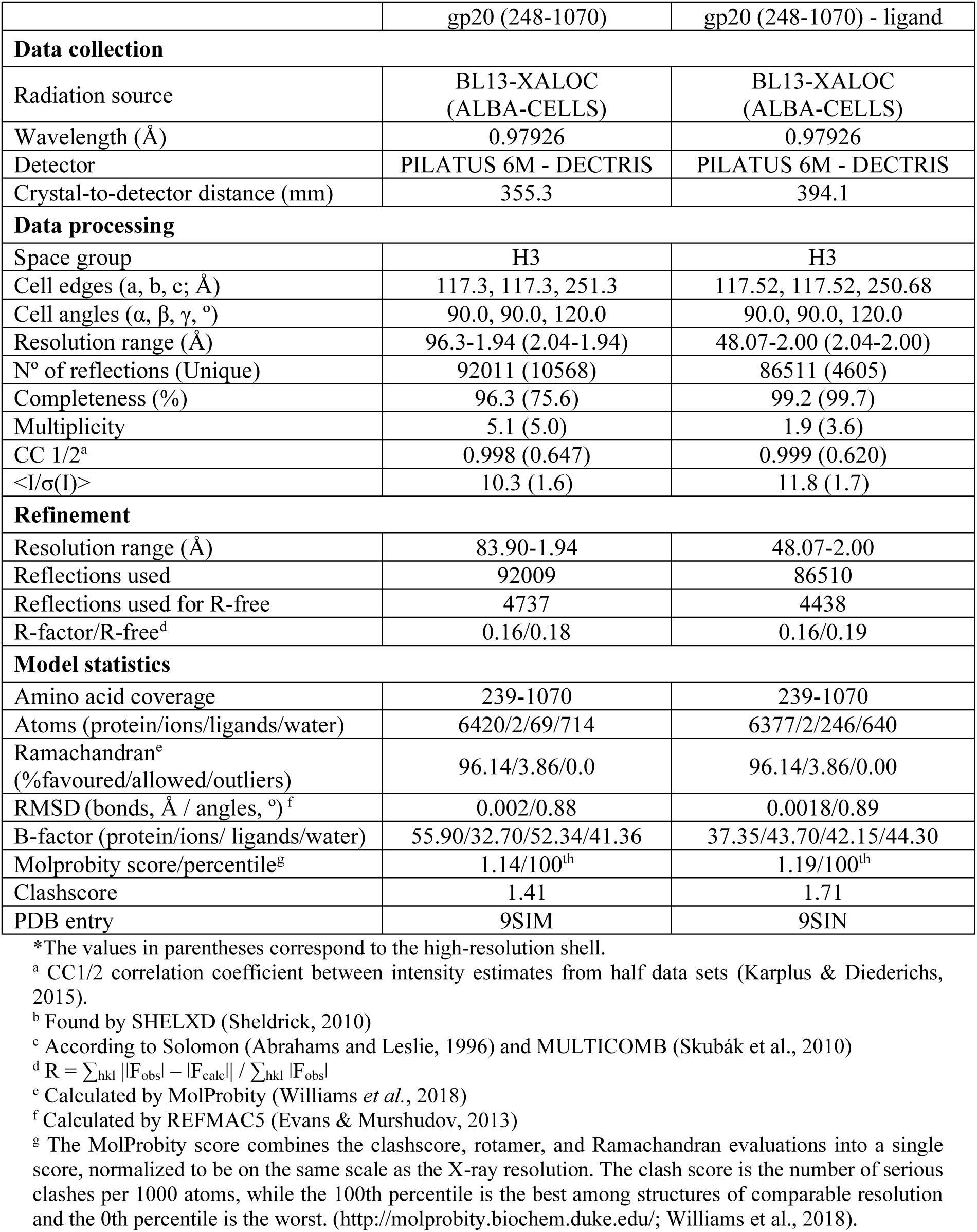
Crystallographic statistics of gp20(248-1070) datasets *The values in parentheses correspond to the high-resolution shell.

**Figure 2.**
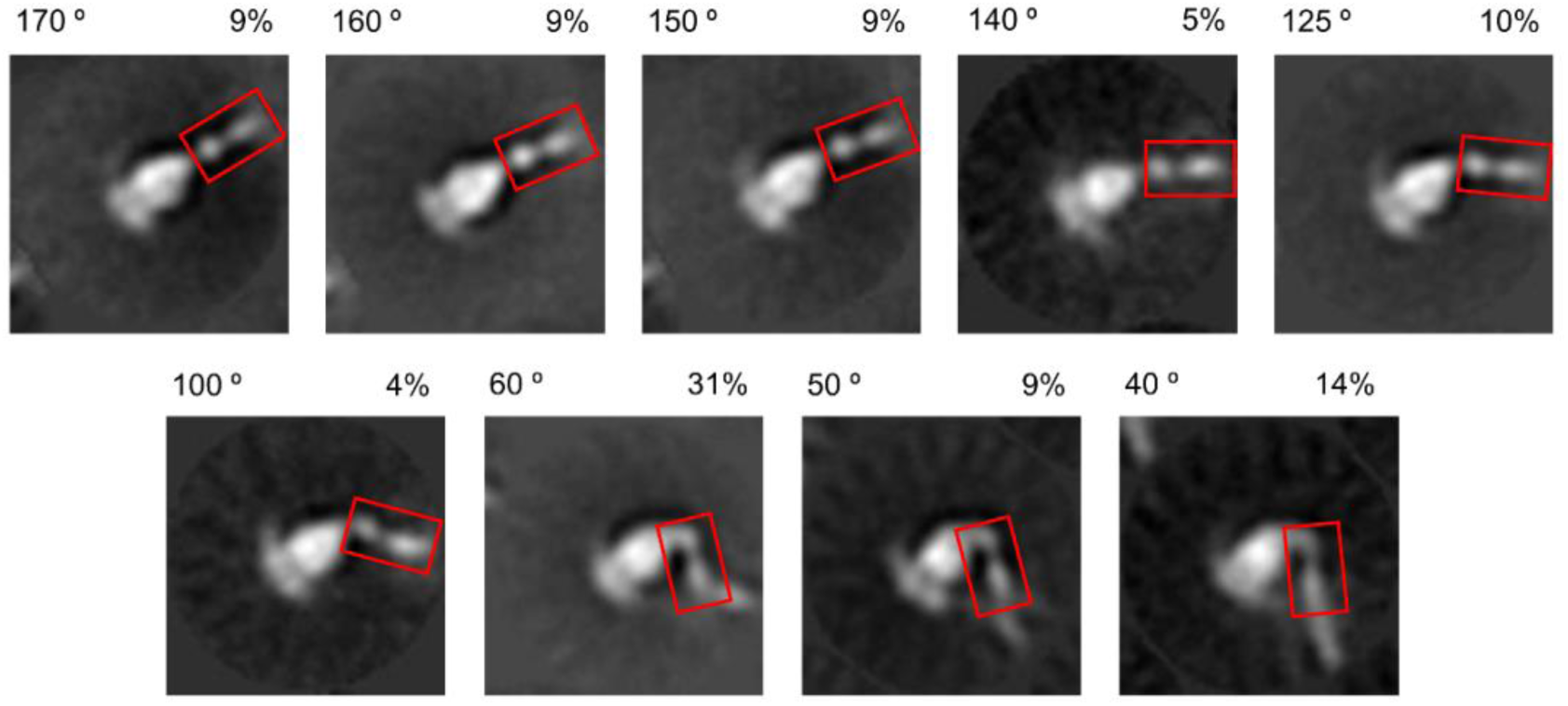
Phage-binding domain flexibility. Nine class averages display the gp20 N-terminal region (highlighted in the red rectangle) bent at different degrees. These averages are derived from nearly 15,000 particles. The approximate angle relative to the barrel and the percentage of particles in each class are indicated above the corresponding images.

The petal domain has a novel fold with no structural homologues. It is composed of two subdomains (Figure 3*g*). The larger subdomain adopts an alpha/beta fold (residues 778-825 and 907-1070), consisting of seven alpha-helices and two 3_10_-helices surrounding a parallel four-stranded beta-sheet. The smaller subdomain forms a six-stranded antiparallel beta-barrel, spanning residues 826-906. The loops between strands 4 and 5 and strands 6 and 7 are extended, creating a groove with the large alpha/beta subdomain.

The alpha/beta subdomain is structurally homologous to members of the SGNH-hydrolase family, which possess esterase activity. The RMSD values for PbAcXE, TesA, IAH1 and OatA proteins, representatives of this family, are 2.4 Å (PDB codes: 7TOI, 4JGG, 3MIL, 6WN9) (Figure S1a). A key feature of this superfamily is the presence of four conserved residues – Ser, Gly, Asn and His – found in conserved blocks I, II, III and V, respectively (Akoh *et al*., 2004). In the petal domain, these conserved residues are Ser787, Gly813, Asn937 and His1038 (Figure 3*f*). Ser787, Asp1035 and His1038 form the putative catalytic triad. These three amino acids are connected by a network of hydrogen bonds. In the apo-petal domain structure, Ser787 adopts a dual conformation, with its hydroxyl oxygen forming hydrogen bridges with the His1038 Nδ1 and the Gly813 amide (Figure 3*f*).

### 3.2. Identification of four O-antigen binding sites per gp20 monomer

To further investigate the receptor-tailspike interaction, we crystallised the two gp20 constructs with fragments of the Anatum O-antigen receptor. Gp20(248-1070) crystals were soaked with hexasaccharide fragments, which have the sequence D-Galp[6Ac]-α-1–6-D-Manp-β-1–4-L-Rhap-α-1–3-D-Galp[6Ac]-α-1–6-D-Manp-β-1–4-L-Rhap-α-1–ROH. The best crystal diffracted at 2.0 Å resolution. The resulting structure revealed three bound hexasaccharides: two located around the negatively charged groove of the beta-helix and one in the beta-sandwich domain. From here on, hexasaccharides will be referred to as Hexa1, Hexa2 and Hexa3 in protein N-to C-termini order (Figure 4*a-d*). Additionally, individual saccharides will be named starting from the non-reducing end galactose (Gal1, Man2, Rha3, etc.).

**Figure 3.**
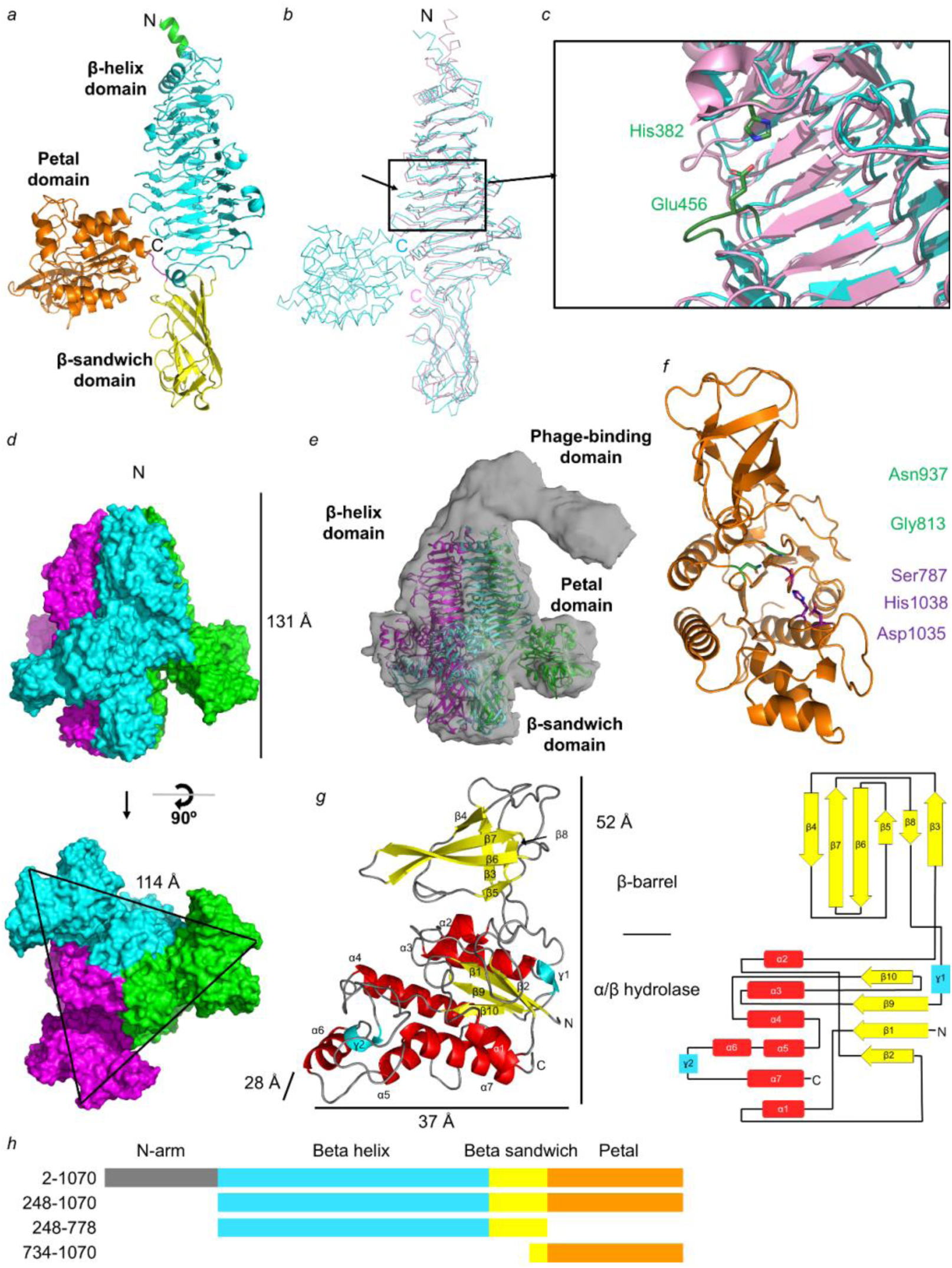
Gp20 crystal structures. (*a*) Ribbon representation of the gp20(248-1070) monomer, with the affinity tag residues in green, the beta-helix domain in cyan, the beta-sandwich domain in yellow, the linker in magenta and the petal domain in orange. (*b*) Wire representation comparing gp20(248-1070) (cyan) and gp208 (light pink), with the N-and C-termini labelled. The black arrow highlights loop 456-459. (*c*) Close-up view of loop 456-459, showing the salt bridge between Glu456 and His382 (*d*) Surface representation of trimer gp20(248-1070), with each chain in a different colour. The side view is above, and the bottom view is below. The overall dimensions of the gp20(248-1070) trimer are 131 x 114 Å (height x triangle side, measured from petal Gly878 to petal Gly878; the furthest residues from the centre, 60 Å away). (*e*) Side view of the epsilon15 tailspike cryo-EM map (13 Å resolution, EMDB entry: EMD-5209, Murata *et al*., 2010) in grey, with the fitted gp20(248-1070) trimer coloured as in (*d*). (*f*) Close-up of the petal domain, highlighting the catalytic residues (Ser787, Asp1035 and His1038) in violet, with the oxyanion hole residues in green. (*g*) Ribbon representation (left) and topology diagram (right) of the C-terminal petal domain, with alpha-helices in red (α), beta-strands in yellow (β), 3_10_-helices in cyan (ɣ), and loops in grey. This fragment is divided into two subdomains: the beta-barrel and the alpha/beta hydrolase fold, forming a structure of about 52 x 38 x 28 Å (height x width x depth). (h) Domain organization of gp20 constructs. Schematic representation showing the N-arm (grey), β-helix (light blue), β-sandwich (yellow), and petal (orange) domains for gp202–1070), gp20(248–1070), gp20(248–778), and gp20(734–1070).

**Figure 4.**
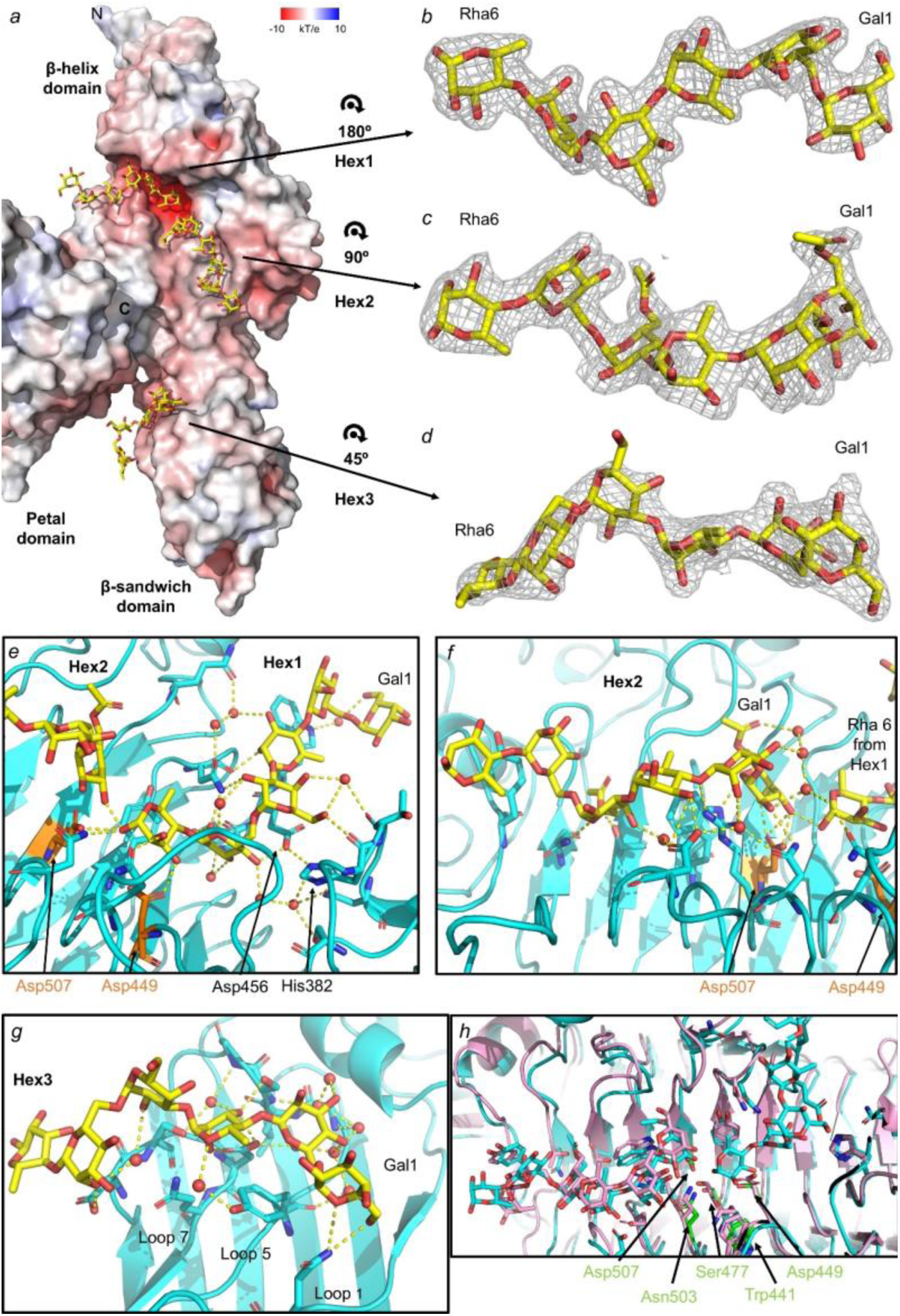
Gp20(248-1070) with three bound hexasaccharides. (*a*) Surface representation of gp20(248-1070) structure coloured by surface charge: red for acidic, blue for basic, and white for neutral. The three bound hexasaccharides are shown in stick representation (yellow and red). Individual depictions of Hexa1 (*b*), Hexa2 (*c*) and Hexa3 (*d*) are presented with their 2Fo-Fc electron density maps contoured at 1σ (grey). Close-up views of Hexa1 (*e*), Hexa2 (*f*) and Hexa3 (*g*) binding sites display the hydrogen bonds (yellow dashed lines). Hexasaccharides are depicted as in (*a*), gp20(248-1070) shown as ribbon (cyan), residues as sticks (cyan, Asp507 and Asp449 in orange) and water molecules as spheres (red). (*h*) Superposition of gp20(248-1070) and gp208 (pink) with their receptors. The gp20(248-1070) residues mutated for the endorhamnosidase assays are coloured green.

Hexa1 binds to the negatively charged patch and the His382-Asp456 bridge in the beta-helix (Figure 4*a*). It interacts with amino acids in parallel beta-sheet 3 of rungs 4, 5, 7 and 8, along with their surrounding loops. The binding involves mainly direct or water-mediated hydrogen bonds connecting with all saccharides, supplemented by some aromatic stacking interactions (Figure 4*e*). Hexa2 also binds to the beta-helix domain, extending from the distal end of the negatively charged patch to the end of the domain (Figure 4*a*). This hexasaccharide occupies a nearly identical position to that observed in the gp208 structure (Figure 4*h*), with binding residues located in the PB3s of rungs 7-12 and adjacents loops. The interactions are predominantly hydrogen bonds, with an aromatic stacking interaction between Tyr630 and Man5, also found in the gp208 structure. Notably, the Gal1 of Hexa2, located on the negatively charged patch, forms the most hydrogen bonds with the protein (Figure 4*f*). The beta-helix groove width ranges from 7.5 to 17.5 Å, measured at Hexa1 Man5 and Hexa2 Man2, respectively, and spans 34 Å long. Finally, Hexa3 binds to loops 1, 5 and 7 of the proximal region of the beta-sandwich domain, which faces the petal domain (Figure 4*g*). Although the soaked hexasaccharide had an acetyl group bound to the C6 of the galactose residues, this group is only resolved in Hexa2 galactoses (Figure 4*c*), likely due to its flexibility or the esterase activity of the petal domain.

Gal1 of Hexa2 and the reducing Rha6 Hexa1 are connected via a hydrogen bond (Figure 4*e,f*), forming an interface just above the negatively charged patch. At this site, the carboxyl oxygens of Asp449 and Asp507 are 5.3 Å apart, indicating a potential catalytic site. According to the literature (Davies & Henrissat, 1995), this distance suggests that gp20(248-1070) uses the retaining mechanism to hydrolyse *Salmonella enterica* serovar Anatum O-antigen polysaccharides.

Petal domain crystals were soaked with nonasaccharide fragments of the Anatum O-antigen. The best crystal diffracted X-rays to 2.08 Å, resolving a pentasaccharide within the petal domain groove (Figure 5*a*). This pentasaccharide lacks the reducing rhamnose found in the hexasaccharides of the gp20(248-1070) structure. The mannose at position 5, now the reducing end, forms only one water-mediated hydrogen bond with the protein. Given its position, the following saccharide (rhamnose) would be too distant from any protein residue to interact. The lack of interaction likely renders the remaining nonasaccharide too flexible to be resolved in the crystal structure. The acetyl groups on the galactoses are also partially unresolved, possibly due to their flexibility or the esterase activity of the petal domain.

**Figure 5.**
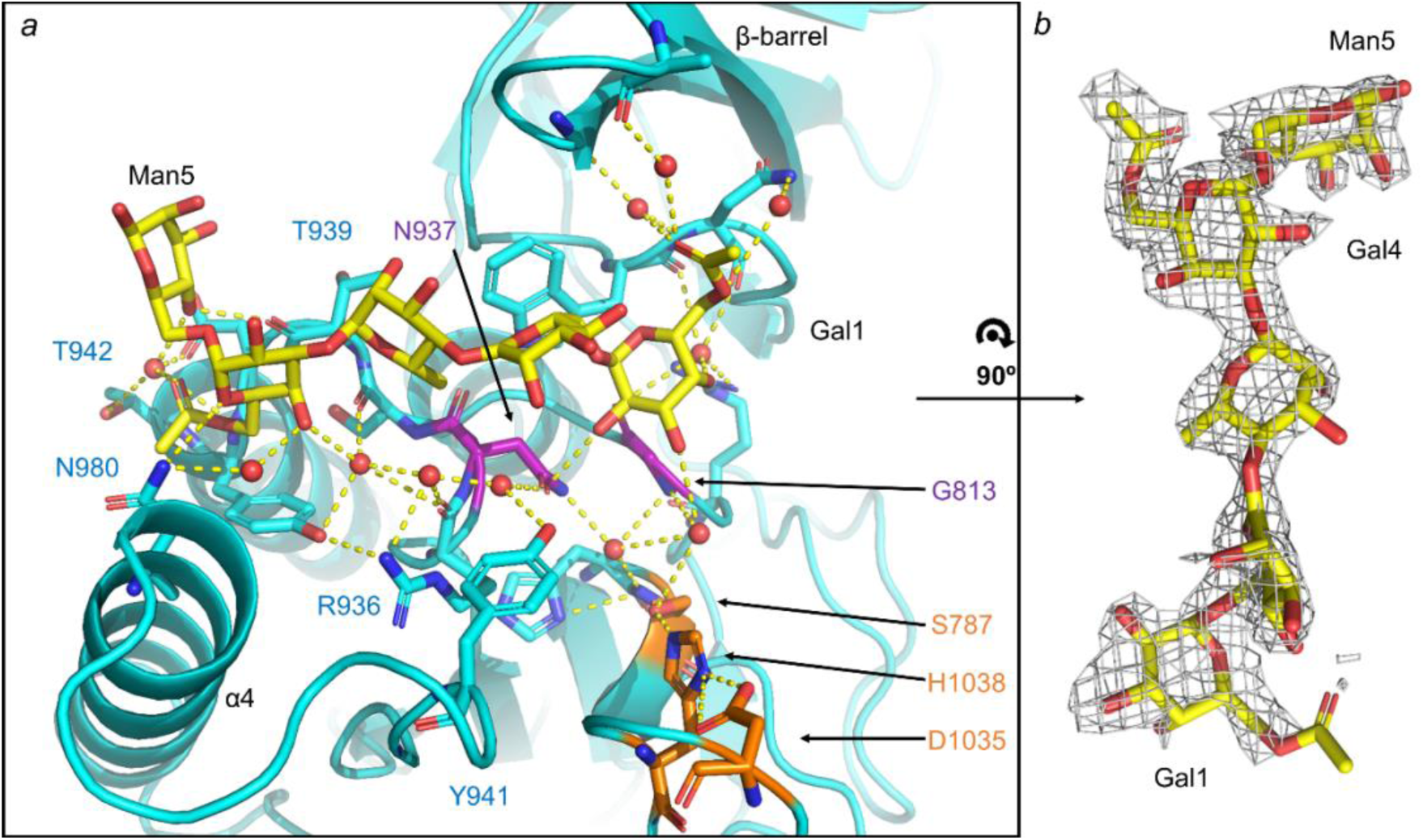
**Petal-oligosaccharide complex.**(*a*) Close-up view of the pentasaccharide and its petal biding site in the petal domain. The petal domain is displayed in ribbon form (cyan), with pentasaccharide-interacting residues depicted as sticks. The putative catalytic triad is highlighted in orange, while the oxyanion hole residues are in purple (with Ser787 present in both groups). Water molecules involved in the interaction network are represented as small red spheres, and hydrogen bonds as yellow dashed lines. (*b*) Stick representation of the pentasaccharide (yellow) with the 2Fo-Fc electron density map contoured at 1σ (grey).

The oligosaccharide interacts with the petal domain, primarily engaging with the alpha/beta subdomain. These interactions are mediated by a network of hydrogen bonds, many of which involve water molecules. The saccharide closest to the esterase site is Gal1, the non-reducing end. However, its ester group is located 12 Å from Ser787. Unlike in the apo-protein, Ser787 adopts a single conformation in this structure.

An Anatum O-antigen fragment is present in the petal domain crystal structure but absent in the petal domain of the gp20(248-1070) crystal structure. This discrepancy arises from the packing arrangement of the gp20(248-1070) tailspikes within the crystal. Each tailspike forms a pyramidal tetrahedron with three others (Figure S2*a*), interacting at four regions: the N-terminal end and the three petal grooves. The N-terminal alpha-helix consists of the first three amino acids from the truncated gene and the final eight amino acids of the His-tag. Each of the three N-terminal helices engages with the groove of a petal domain from a different tailspike (Figure S2*b*). The oligosaccharide bound to the petal domain structure occupies the same position (Figure S2*c*). Consequently, the hexasaccharides added during the soaking experiments were unable to reach their binding site.

### 3.3. Key residues of gp20 mediate endorhamnosidase activity via an inverting hydrolysis mechanism

Several single-point, mutant variants of gp20(248-1070) were created in order to investigate the possible roles of specific amino acids positioned either near the Hexa1 Rha6 and Hexa2 Gal1 interface (D449N, S477A, N503D and D507N) or within the petal triad (S787A, H1038A) on the esterase estendorhamnosidase activities (Figure 4*h*, 6*a-b*). A seventh mutant, W441R, was createdbecause of earlier observations showing that this gene 20 mutation enables E15 phage to infect both *S. enterica* serovar Anatum A1 cells (displaying antigen O10) and *S. enterica* serovar Anatum A1(E15) cells (displaying antigen O15), with equal efficiency (McConnell lab, unpublished data).

The hydrolytic activity of the WT fragments and their mutants on the Rha-alpha (1–3)-Gal linkage was evaluated using the MBTH assay and by NMR spectrometry (Figure 6*c-d*, S5*a*, S6*a*, S7*a*, S8*a-c*, S9*a-b*, S10*a-b*), both explained in the Materials and Methods section. Briefly, in the MBTH assay, the reaction of two MBTH molecules with reducing ends in the media produces a blue colour with peak absorption at 655 nm. In the NMR approach, first the polysaccharide structure was analysed by acquiring a set one-and two-dimensional NMR spectra (1D-^1^H and 2D-TOCSY, HSQC and DOSY, Figure S4) confirming that its structure corresponds to the O-antigen 10 as described previously (Gajdus *et al*., 2008). Next, the enzymatic reactions were directly followed in the spectrometer by monitoring changes in signal intensities of the anomeric hydrogen of newly formed reducing-end rhamnose residues. This approach enabled us to track the reaction dynamics under different conditions. Additionally, DOSY experiments were recorded at the start, at selected intermediate points, and at the end of the reaction to observe the decrease in the molecular size of the polysaccharide, resulting from the cleavage by the endoglycosidase activity (Figure S5*c*, S6*e*, S7*e*).

**Figure 6.**
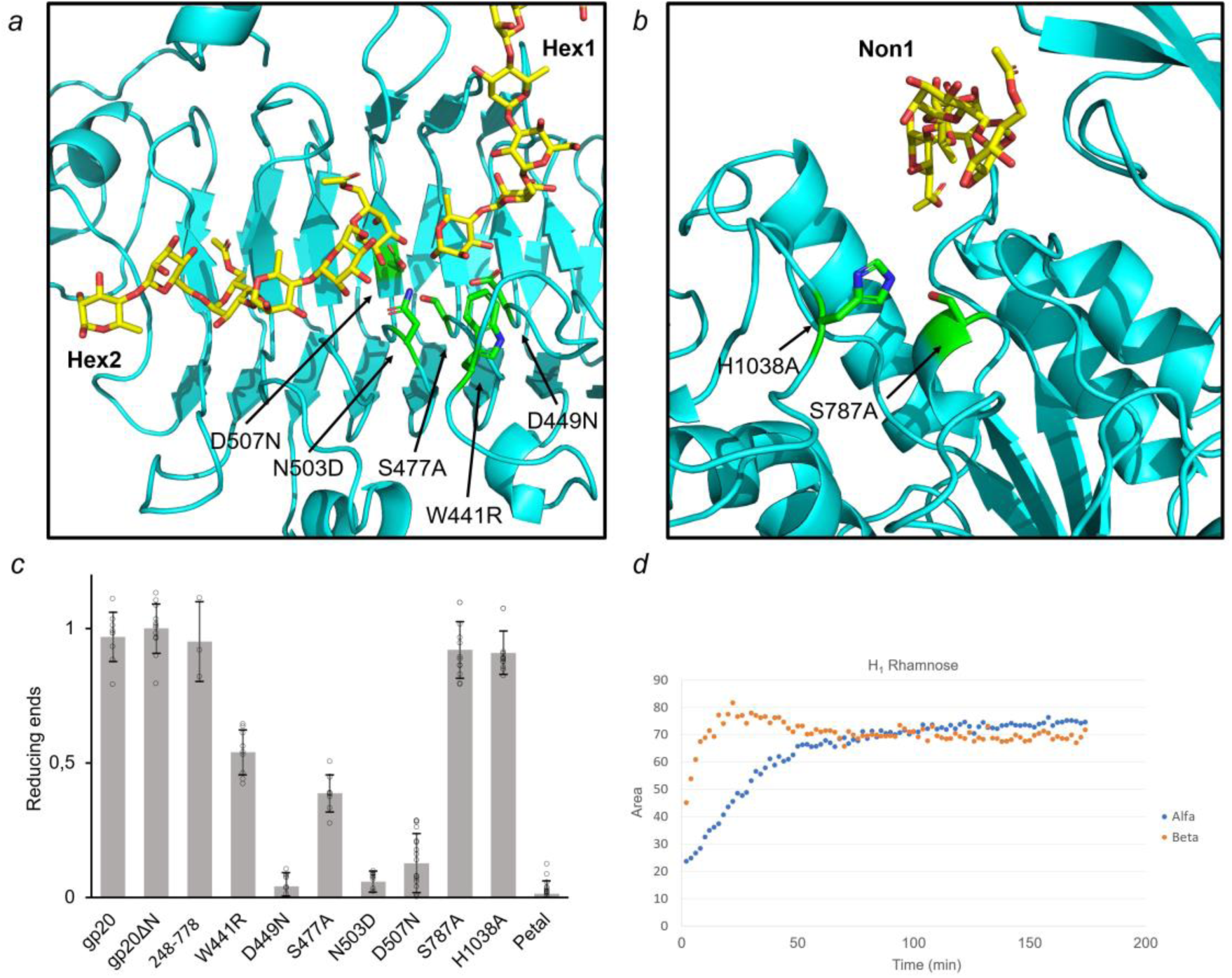
**Endorhamnosidase activity.**(*a*) Putative endorhamnosidase and (*b*) esterase sites, with gp20 shown in cyan and mutated amino acids in green. (*c*) Relative amount of reducing ends produced by gp20 constructs and single-point mutants (1 µM), where the non-hydrolysed polysaccharide signal is set as 0.0 and the signal from gp20(248-1070)-hydrolysed polysaccharide is set as 1.0. Statistical comparisons were performed with a Student’s t-test (n ≥ 3) (*d*) A plot over time of the integrals of the peaks corresponding to α and β anomeric protons of the reducing-end rhamnose shown in Fig. S5d, generated during the incubation with gp20(248-1070) (1 µM). The data were obtained from NMR spectra recorded over the course of the incubation period. The y-axis corresponding integrated area of NMR peaks has arbitrary units.

Reducing end production in reactions containing the petal domain did not differ statistically from negative control reactions involving the polysaccharide alone (Figure 6*c*), indicating that its putative active site is not involved in O-antigen cleavage and can therefore serve as a negative control. On the contrary, full-length gp20, gp20(248-1070) and gp20(248-778) proteins all produced new reducing ends at a similarly high efficiency, and exhibited similar results generating new reducing ends leading to the use gp20(248-1070) as a positive activity reference for all of the mutants that had been created from. The mutants D449N, N503D and D507N showed the lowest production of reducing ends, statistically comparable to the petal domain, indicating their likely involvement in the O-antigen cleavage reaction (Figure 6*c*, 0). In contrast, esterase mutants S787A and H1038A had activity levels similar to gp20(248-1070), while W441R and S477A displayed intermediate activities, suggesting these residues are important for oligosaccharide binding but likely not directly involved in catalysis.

The NMR analysis of the gp20 constructs confirmed the MBTH assay results, corroborating the presence of endorhamnosidase activity in gp20(248-1070) (Figure 6*d*, S5*a,b,d*, S7*a,b,d*) and the absence in the petal domain (Figure S6*a,e*), on the Anatum polysaccharide (0). The key mutants D449N, N503D and D507N demonstrated their lack of activity on the polysaccharide (Figure S8*a-c*). The gp20(248-778) construct exhibited endorhamnosidase activity (Figure S7*a*), further confirming that the petal domain does not influence its activity. Notably, NMR revealed that gp20(248-1070) and gp20(248-778) employ an inverting mechanism during hydrolysis, converting the alpha configuration of rhamnose in the polysaccharide to beta configuration in the new reducing ends generated in the reaction (Figure 6*d*, S5*d-f*, S7*b-d*), which then mutarotates nonenzymatically until reaching the alpha/beta anomeric equilibrium of 53% alpha and 47% beta (Figure 6*d*).

### 3.4. Esterase activity of the Petal domain on galactose acetyl groups

The structural similarity of the alpha/beta subdomain to proteins in the SGNH-hydrolase family led us to hypothesise that the petal domain might exhibit esterase activity (Figure S1*a*). We therefore used NMR to measure the esterase activity of the petal domain alone, gp20(734-1070), as part of the gp20(248-1070) construct and in the absence of the petal domain with the gp20(248-778) construct. Only the constructs with the petal domain displayed esterase activities at comparable levels (Figure 7*a-c*, S5*g*, S6*c*, Table 4), suggesting the esterase activity depends only on the petal domain.

**Figure 7.**
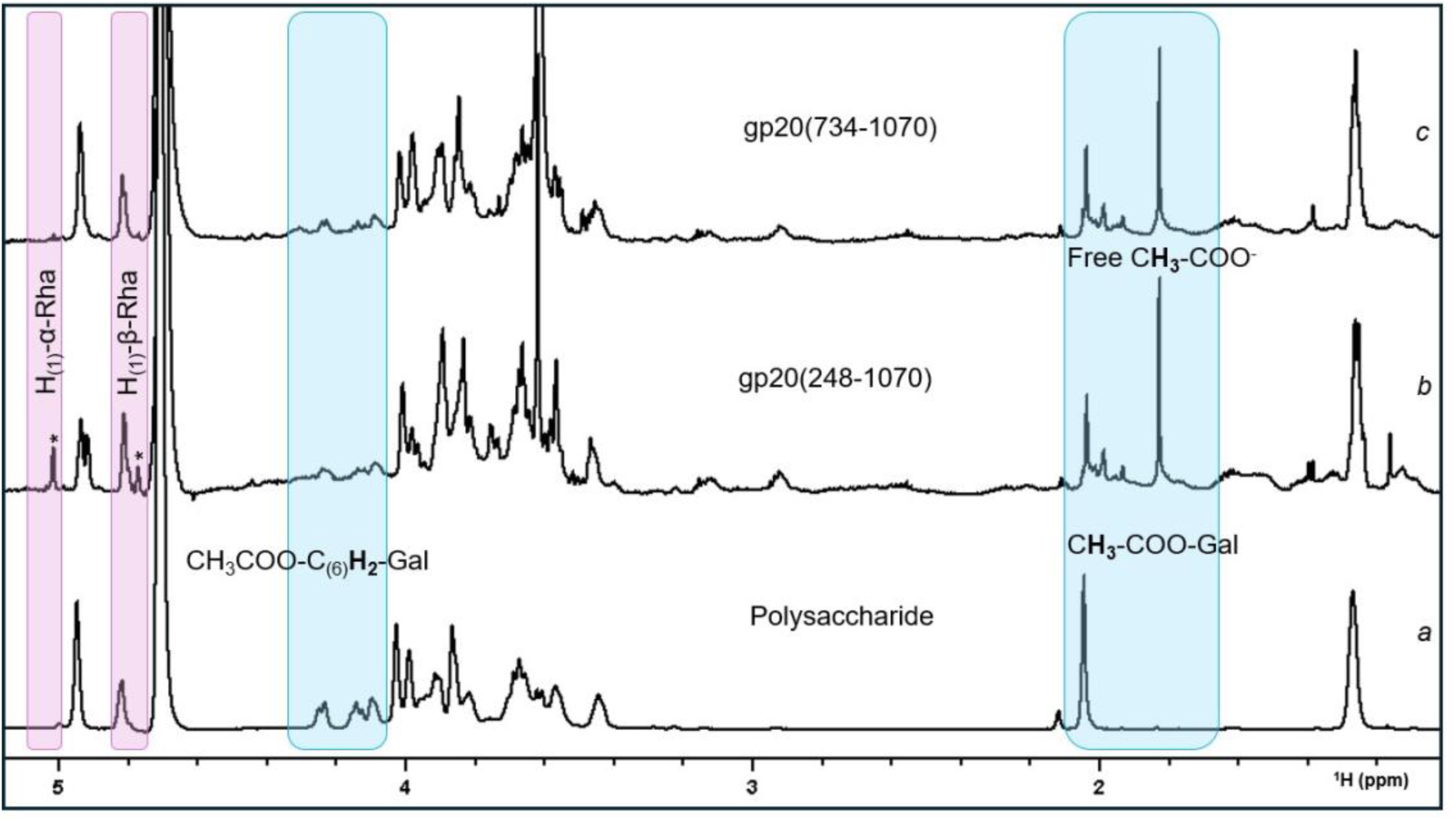
**Esterase activity.**^1^H NMR spectra of: *(a)* polysaccharide in the absence of any protein; *(b)* after 24 h in the presence of gp20(248-1070) and *(c)* in the presence of gp20(734-1070) in ratios protein:ligand 1:50. Highlighted are the changes due to esterase activity, appearance of free acetate and disappearance of signals from CH_2_ at position 6 of acetylated galactose (blue) and endorhamnosidase activity, appearance of α and β anomeric protons (*) of rhamnose at the new reducing ends (pink).

**Table 4.** Endorhamnosidase activity and esterase activity measured by NMR

| Gp20 constructs and mutants | Endorhamnosidase activity | Esterase activity |
| --- | --- | --- |
| Gp20(248-1070) | + | + |
| Petal | - | + |
| Gp20(248-778) | + | - |
| Gp20(248-1070) D449N | - | + |
| Gp20(248-1070) N503D | - | + |
| Gp20(248-1070) D507N | - | + |
| Gp20(248-1070) S787A | + | - |
| Gp20(248-1070) H1038A | + | - |
| Petal S787A | - | - |
| Petal H1038A | - | - |

To further investigate this potential activity, we mutated the key residues of the SGNH-hydrolase family esterase site in the petal domain, S787A and H1038A in gp20(248-1070) and in gp20(734-1070) (Figure 6*b*). After 24 h, the signals for the protons of the methyl group (signal at 2 ppm) and the exocyclic CH_2_ (signals 4-4.3 ppm) of acetylated galactose remained unchanged in the spectra (Figure S9*a-b*, S10*a-b*), confirming that the petal domain possesses esterase activity against the acetyl group of the galactoses (Figure S5*i*).

## 4. Discussion

The epsilon15 tailspike is comprised of four distinct regions; at the N-terminus is a phage binding domain, followed by a beta-helix domain, a beta-sandwich domain and petal domain (Figure 3*a*). Because the phage-binding domain makes the tailspike too flexible for crystallisation, we chose to study it using microscopy methods. This work revealed a phage-binding structure comprised of two knobs connected by a shaft, with a shorter and more flexible shaft connecting the two knobs to the beta helix domain. AlphaFold3 analysis (Abramson *et al*., 2024) predicted a similar arrangement of motifs in the phage-binding arm (Figure S*4a*), with the flexible shaft bending at angles of approximately 170 ° (in one case) and 60 ° (in four others), aligning with the microscopy data (Figure 2).

Advanced protein prediction methods, such as Alphafold3, were unavailable when our structural studies were finished. When applied to gp20, Alphafold3 gives results closely matching our experimental structure for the petal domain (RMSD = 0.4-0.5) as well as for the combined beta-helix and beta-sandwich domains (RMSD = 0.6-0.7), structurally homologous to Det7 gp208 tailspike. As for the gp20(248-1070) fragment containing the beta helix, beta-sandwich and petal domains, the RMSD value was less impressive (2.8 to 10 Å^2^), probably because of the petal domain can be oriented in more than one way relative to the main body (Figure S3*b*).

The function of the three C-terminal domains of the E15 tailspike is to bind to the *S. enterica* serovar Anatum O-polysaccharide. The reducing ends of the crystal-bound Hexa1, Hexa2 and Hexa3 oligosaccharides all point towards the C-terminal, distal end of the tailspike; i,e, towards the lipid A anchor of the lipopolysaccharide (LPS) located at the outer membrane surface. This is also true for the hexasaccharide present in the Det7 gp208 structure. The common orientations of all these oligosaccharides argue against them being artefacts of crystallisation. The positioning of the downstream end of Hexa1 next to the upstream end of Hexa2 in the vicinity of amino acids D449, N503 and D507 is consistent with our experimental evidence showing that these three amino acids comprise the endorhamnosidase catalytic site. The 23.7 Å distance between Rha6 O1 of Hexa2 and Gal1 O3 of Hexa3 fits nicely within the 22.1 to 24.5 Å range calculated for modelled hexsaccharides (Figure 4*a*). Hence, another hexasaccharide could easily fit in between Hexa2 and Hexa3. Since the bacterial polysaccharide consists of chains with varying numbers of repeating units, we hypothesised that the tailspike protein would cleave the polysaccharide into fragments compatible with its binding pockets. Specifically, we expected a single fragment to occupy the three binding pockets within the β-sandwich and β-helix domains. However, our attempts to crystallise gp20(248-1070) with whole Anatum polysaccharide were unsuccessful.

The petal domain displays both oligosaccharide binding and esterase activities (Figure 5*a*, 7), with both functions residing within a groove that borders the beta-barrel subdomain and the alpha/beta hydrolase domain. Surprisingly, Ser787, which is the conserved catalytic residue in the SGNH-hydrolase family, lies 12 Å away from the nearest ester linkage connecting galactose with an acetyl group. If some oligosaccharide flexibility is required to bring the ester linkage close to the catalytic amino acids (Figure 5*a*), it could help explain the relatively slow kinetics of esterase activity (Figure S6*d*), relative to the endorhamnosidase activity (Figure S5*a*). Other structurally similar esterases show similarly slow cleavage efficiencies, like OatA on N-acetylmuramic acid (Jones *et al.,* 2020, PDB code: 6WN9) or PbeAcXE on xylans (Penttinen *et al*., 2022, PDB code: 7TOI), while the tailspike gp63.1 of phage G7C releases all the acetyl group from the bacterial receptor in 2 h (Prokhorov *et al*., 2017).

Recent work has revealed that SGNH hydrolase domains are prevalent among receptor-binding proteins (RBPs) in temperate *Klebsiella pneumoniae* phages, often surpassing the classical pectin lyase-like depolymerase domains in frequency. These SGNH domains are proposed to mediate capsule deacetylation, a mechanism distinct from depolymerisation, which may facilitate phage infection by modifying host polysaccharide surfaces (Otwinowska *et al*., 2025)

The ability of gp20(248-778) to fold without the petal domain is surprising since the traditional understanding of tailspikes is that they commence their folding from the C-terminal end (Gage & Robinson, 2009), and thus, C-terminal modifications might hinder correct folding. This special feature may make gp20 an interesting subject for biotechnological applications. Trimeric tailspikes are known for their thermal stability (Barbirz *et al*., 2009). Hence, a different enzyme could be attached to the C-terminal end of gp20(248-778) instead of the petal domain to obtain a stable complex, either in the isolated tailspike or in a genetically modified bacteriophage.

Early studies (Hagiwara *et al*., 1966; Wright, 1971) showed that acetyl groups are not a requirement for infection of *S. enterica* serovar Anatum by epsilon15 phage. The opposite is apparently true for phage G7C (Prokhorov *et al*., 2017). Previous studies have shown that polysaccharide acetylation prevents hydrolysis by rhamnogalacturonases (Kofod *et al*., 1994; Schols *et al*., 1990; Searle-van Leeuwen *et al.,* 1992), inhibits lysozyme penetration into Gram-negative bacteria (Kulikov *et al*., 2017) and acts as a non-specific shield against phage infection (Golomidova *et al*., 2016; Knirel et al., 2015). It is thought that acetylation may facilitate hydrogen bridge formation between LPS chains, thereby slowing down phage progression through the O-polysaccharide barrier. Our in vitro results show that endorhamnosidase activity is faster than esterase activity, reducing esterase to a side role that, in the case of E15, is apparently dispensable. It is not yet known if the tailspike endorhamnosidase displays differing levels of activity towards Rha-alpha(1-3)-Gal substrates with and without acetylated galactose. Further research is needed to unravel the function(s) of the acetyl group.While this study was in progress, the structure of the Dettilon tailspike was solved (Broeker *et al*., 2019), revealing a central region (residues 262-797) with 43% identity to the beta-helix and beta-sandwich domains (residues 232-772) of the E15 tailspike (Figure 3b). Beta-helix domains featuring negatively charged grooves are present in the tailspikes of bacteriophages P22 (PDB entry: 2XC1; Seul *et al*., 2014), CBA120 (PDB entries: 6NW9, 6W4Q; Greenfield *et al*., 2019, Greenfield *et al*., 2020), K5 (PDB entry: 2X3H; Thompson *et al*., 2010), and SF6 (PDB entry: 2VBK; Müller *et al*., 2008). All of these domains participate in saccharide binding (Davies & Henrissat, 1995, Plattner *et al*., 2017, Dunstan *et al*., 2021) and facilitate endorhamnosidase activity, as observed in P22 (Andres *et al*., 2010; Baxa *et al*., 1996), Det7 gp208 (Broeker *et al*., 2019), Sf6 (Müller *et al*., 2008), and CBA120 TSP2 (Plattner *et al*., 2019). In epsilon15, mutations of Asp449, Asn503, and Asp507 all abolished enzymatic activity and Asp449 and Asp507, located 5.3 Å apart, resemble the typical catalytic dyad seen in retaining glycosidases (Davies & Henrissat, 1995; Zechel & Withers, 2000). That said, according to the CAZY data base (http://www.cazy.org/) (Drula *et al*., 2022), gp20 can be assigned to Glycosyl Hydrolase family 90 (GH90) along with the glycosidases of phages P22 and Det7, for which an inverting mechanism has been proposed but not yet demonstrated experimentally. Interestingly, our NMR studies revealed that epsilon15 follows an inverting mechanism (Figure 6*d* & S5*d*). The question of a retaining versus an inverting mechanism may be explained by assuming that there is some flexibility in the protein, as other works have suggested (Kang *et al*., 2016). Asp449 is situated in a loop between PB2 and PB3 of the sixth rung, so it may be that loop flexibility separates Asp449 from Asp507 during catalysis, creating space for a water molecule to hydrolyse the glycosidic bond via the inverting mechanism. Subsequent mutarotation then would restore the alpha configuration (Figure S5d*,f*).

To our knowledge, this is the only tailspike with four different binding sites for its receptor (Figure 8). The closest example is G7C gp36.1 (Prokhorov *et al*., 2017), where one O-antigen binding site was identified in its crystal structure, and two additional carbohydrate-binding sites were predicted by structural homology. Furthermore, if our model depicted in Figure 8 is accurate, this would be the only known tailspike which binds more than one LPS chain per monomer.

**Figure 8.**
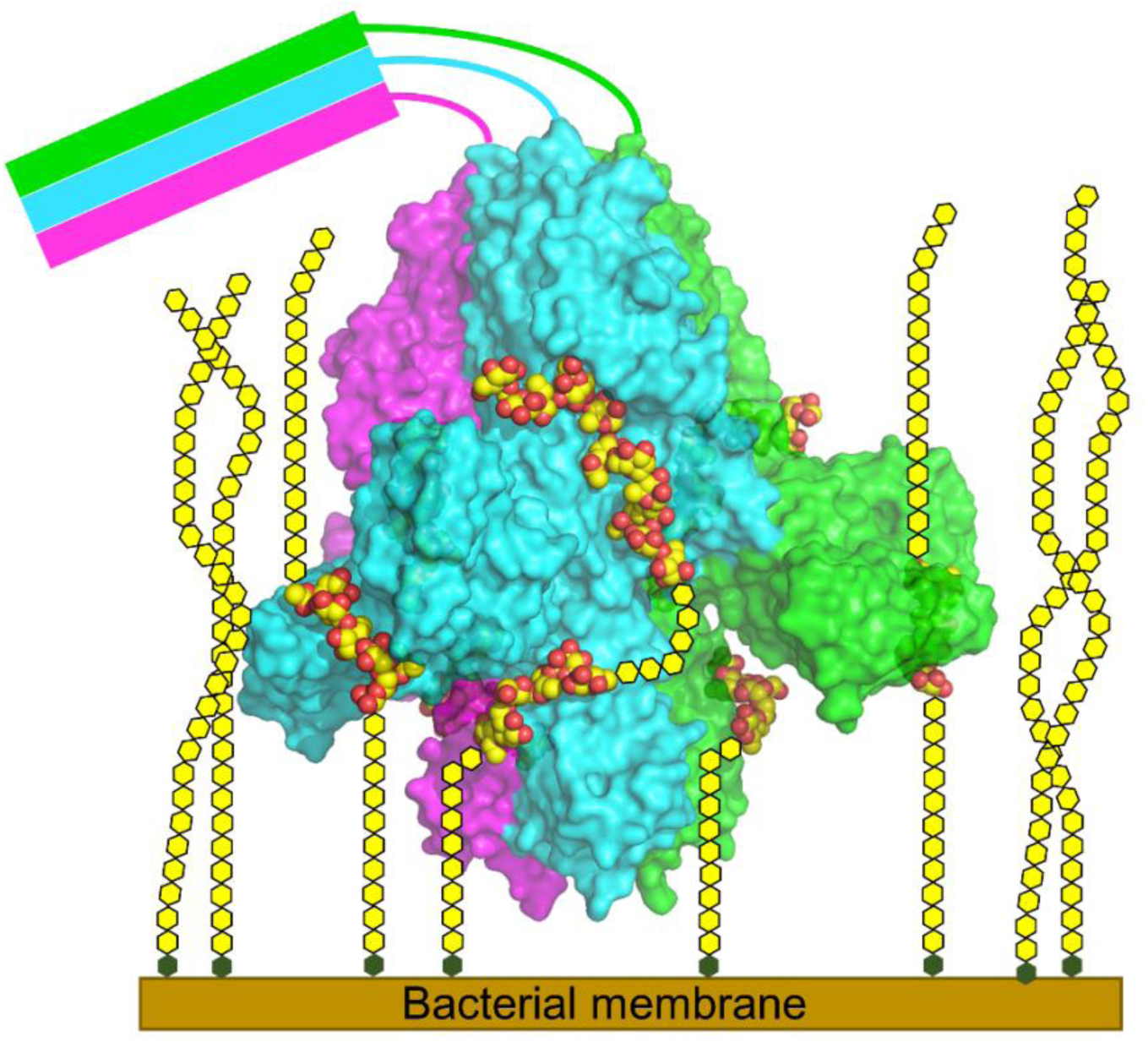
Model of gp20 binding to the bacterial envelope. Surface representation of an epsilon15 tailspike with bound O-antigen fragments, shown in sphere representation. The gp20 monomers are coloured in magenta, cyan and green, while the O-antigen oligosaccharide molecules are depicted with yellow carbons and red oxygens. According to our hypothesis, the oligosaccharides are connected by schematic yellow hexagons: between Hexa2 and Hexa3, between Hexa3 and the membrane, and between the petal pentasaccharide and the membrane. The unresolved N-terminal phage-binding arm is shown schematically.

With the rise of bacterial strains that are multi-resistant and even pan-resistant against clinically used antibiotics (https://www.who.int/health-topics/antimicrobial-resistance; WHO, 2025), phage therapy and phage protein therapy become more important, and with it, a detailed understanding of how bacteriophages recognise and infect their hosts. The high-resolution structure of the *Salmonella* phage epsilon15 tailspike reported here reveals three domains that bind LPS. The C-terminal esterase domain removes acetyl groups, which may make the LPS layer more fluid and allow better phage access. The lectin domain binds but does not modify LPS and is probably important for keeping the phage bound to the host, while the beta-helix domain hydrolyses LPS and allow the phage to move towards the membrane.

## Supporting information

All supplemental figures and tables

## 5.#Acknowledgements

We thank Mariano Marletta, Mara Laguna and Pilar Sánchez-Soriano for technical help in the laboratory. Crystallographic data was collected at the ALBA-CELLS beamline BM13 (XALOC) and beamline staff are thanked for providing excellent crystallographic data-collection facilities. The NMR service from the CIB Margarita Salas (CSIC) is also acknowledged. The MJvR lab was supported by grants BFU2008-01588, BFU2011-24843, BFU2014-53425-P, BFU2017-82207-P and PID2021-125597NB-I00 financed by MCIN/AEI/10.13039/501100011033 and the European Union Next Generation EU/PRTR and FEDER. The Severo Ochoa programme is acknowledged for funding to the CNB-CSIC (SEV-2013-0347, SEV-2017-0712 and CEX2023-001386-S/MICIU/AEI/10.13039/501100011033). MSB was the recipient of FPI predoctoral fellowship associated with grant BFU2014-53425-P, which included funds for a three-month stay in Barbirz lab. FJC acknowledges the financial support of the projects PID2021-123781OB-C22 financed by MICIU/AEI/10.13039/501100011033/ (Spain), P2022/BMD-7278 financed by Madrid Community, Spain and CIBERES, an initiative from the Spanish Institute of Health Carlos III. SB is supported by grants of the German Science Foundation (DFG) BA4046/6-1, BA4046/7-1 and BA4046/8-1. NKB is supported by DFG grants BR5859/2-1 and BR5859/3-1.

