## Supplementary material for "The structure of the *Salmonella* phage epsilon15 tailspike reveals multiple O-antigen binding sites and a protruding esterase domain": All supplemental figures and tables

### 7. Supplementary information

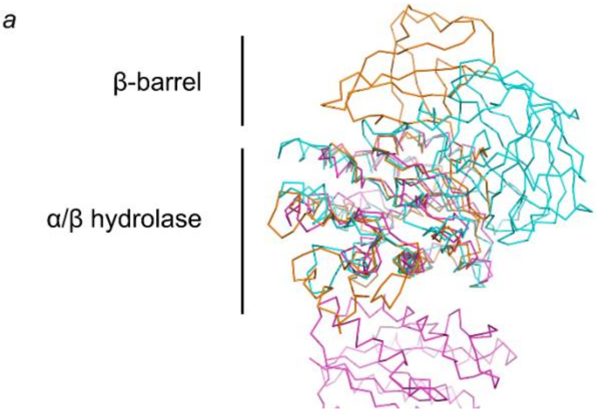


**Figure S1. SGNH homologs of the petal domain.**

*(a)* Wire representation of the petal domain (orange) and the homologous PbAcXE (cyan, PDB code: 7TOI) and TesA (magenta, PDB code: 4JGG) from the SGNH family. All of them shared the β-sheet and the surrounding α-helices typical of the family SGNH.

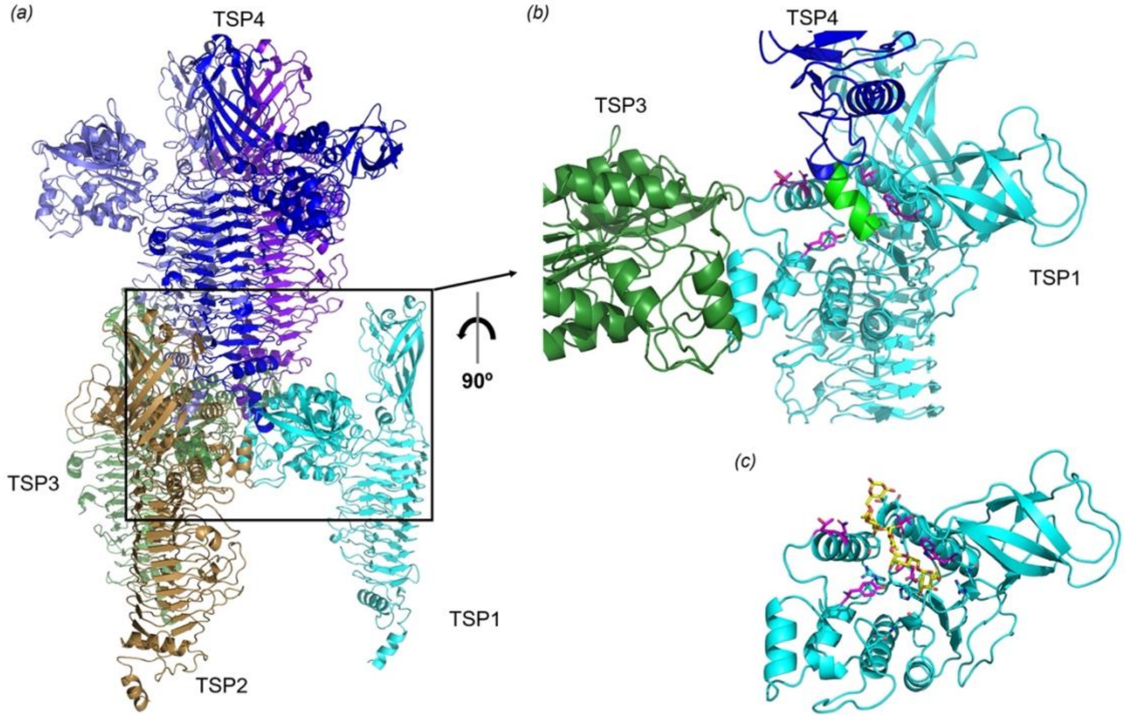


**Figure S2. Gp20(248-1070) crystal packing.**

*(a)* Ribbon representation of four gp20(248-1070) tailspikes in their crystal packing. For the lower TSPs, only the chain contacting TSP4 is shown. Each tailspike is shown in one colour and, for the TSP4, each chain has a different tone of blue. *(b)* A close-up view of the region indicated by a rectangle in *(a)*. For clarity, TSP2 was removed. It would be between the TSP1 and the viewer. The His-tag of TSP4 chain is coloured green. *(c)* Petal domain structure with its bound oligosaccharide. Amino acids contacting the TSP4 N-terminal end in *(b)* and the oligosaccharide in *(c)* are depicted in magenta.

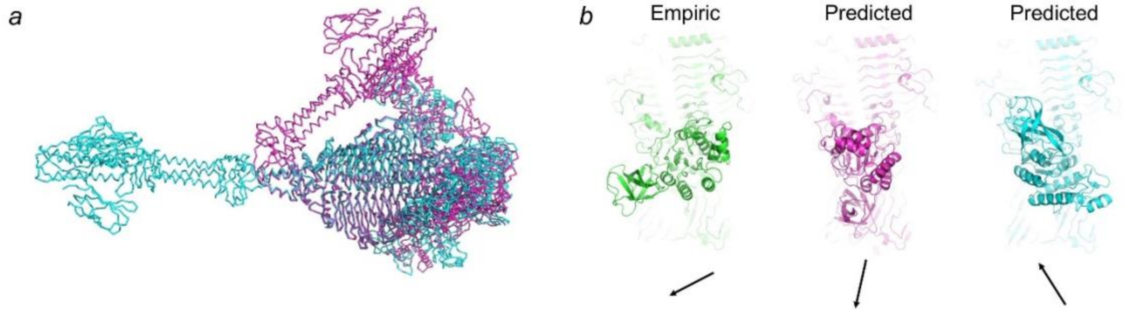


**Figure S3. Structure prediction of gp20.**

(a) Wire representation of two AlphaFold3 (Abramson et al., 2024) predictions of the gp20 trimer (cyan and magenta). (b) Ribbon representation of the experimentally determined gp20 structure alongside the two predicted structures from panel (a). The petal domain is shown in the foreground, while the remaining parts of the structures are partially hidden by the fog. Arrows indicate the orientation of the petal domains, with the arrowhead being in the barrel-subdomain side.

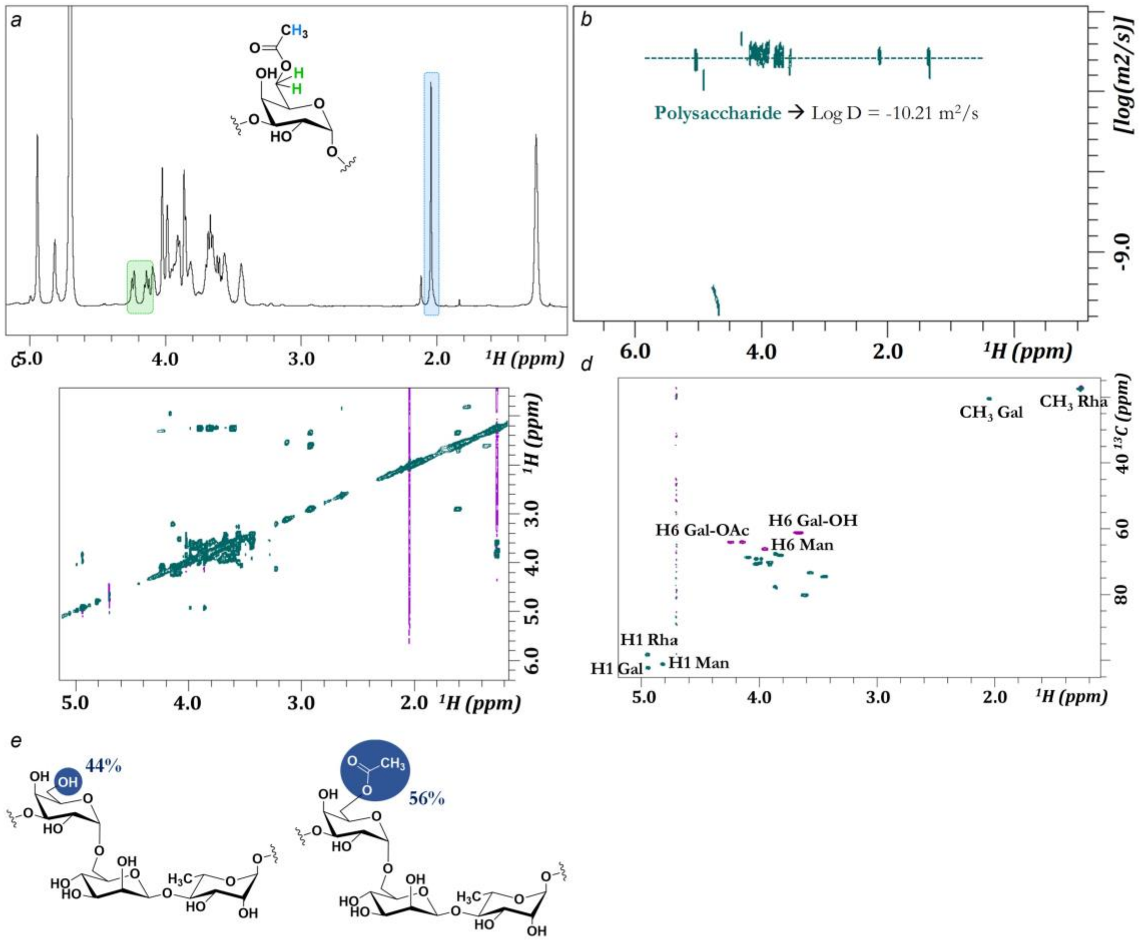


**Figure S4. Polysaccharide analysis.**

*(a)* $^{1}$H-NMR spectrum of the polysaccharide. The highlighted areas correspond to the acetylated galactose moiety, indicating that the polysaccharide was acetylated. *(b)* DOSY spectra of the polysaccharide. The ordinate axis corresponds to the translational diffusion coefficient on a logarithmic scale from higher values (corresponding to smaller-sized molecules) in the lower part to smaller values (larger molecular size) in the upper part. *(c)* $^{2}$D $^{1}$H-$^{1}$H TOCSY NMR spectrum of the polysaccharide. *(d)* $^{1}$H-$^{13}$C HSQC multiplicity edited spectrum (blue/purple). The protons of the $CH_2$ peaks of mannose and of acetylated and non-

acetylated galactose are marked in purple. *(e)* Percentage of acetylated and non-acetylated galactoses obtained from the ratio of acetyl methyl signal of acetylated galactose at 2.05 ppm and the exocyclic methyl signal of rhamnose residues at 1.1 ppm taken as 100% reference.

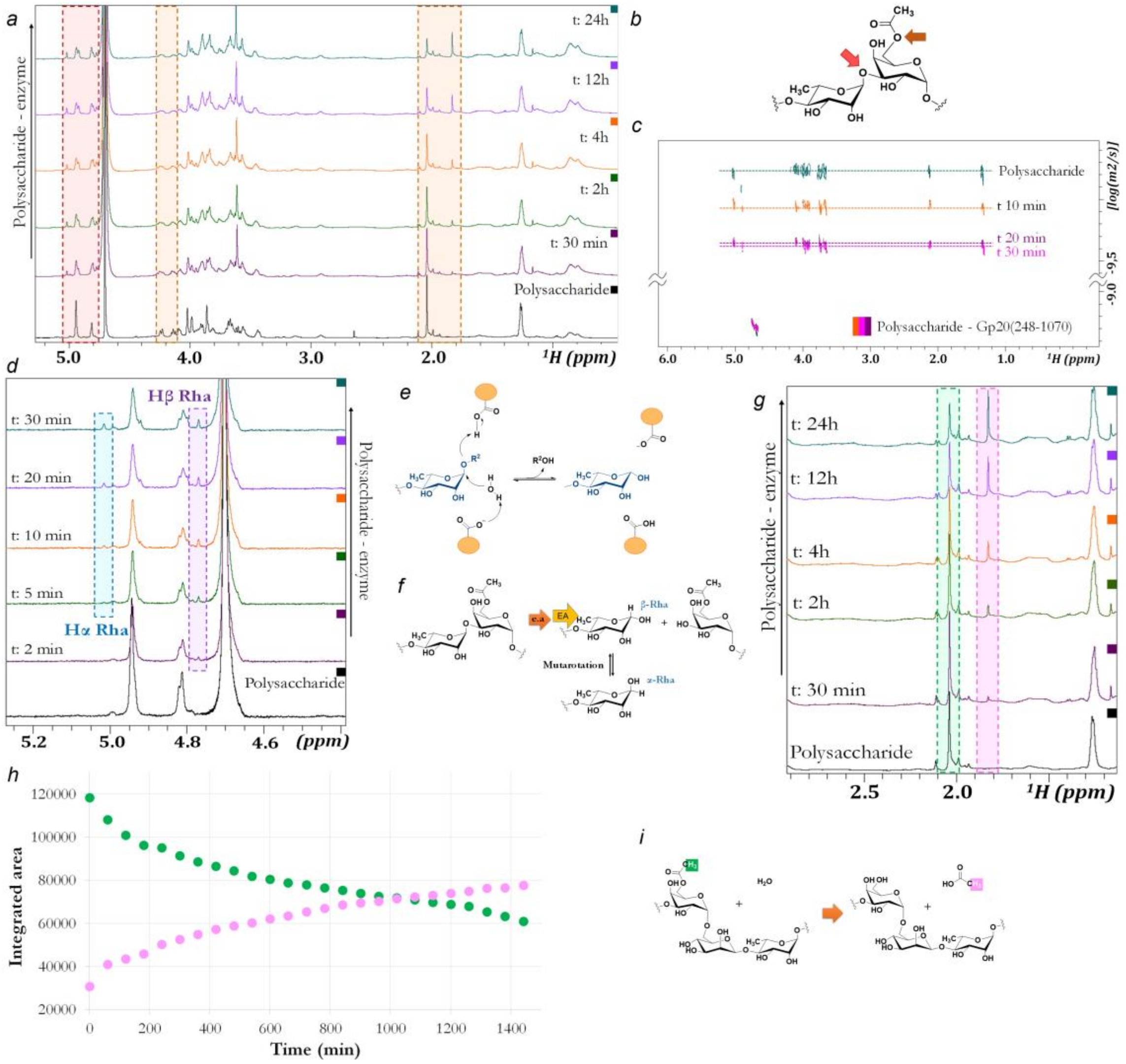


### Figure S5. Enzymatic activity of gp20(248-1070) on the Anatum polysaccharide

*(a)* $^1$H-NMR spectra of polysaccharide in the presence of gp20(248-1070) at different times. The dashed rectangles mark the regions that change over time – red for the endorhamnosidase activity and brown for the esterase activity. The gp20(248-1070) concentration was at 1 µM and the polysaccharide 50 µM (2000 µN of trisaccharide considering a mean value of 40 repeating units per polysaccharide chain). *(b)* Positions on the polysaccharide affected by enzyme activities. *(c)* DOSY spectra of the comparison of polysaccharide size in the absence and presence of enzyme at different reaction times. *(d)* Close-up of $^1$H-NMR spectra from *(a)* at different times of polysaccharide in the presence of gp20(248-1070). The α-, β-rhamnose anomers are shown in boxes, the α-anomer is blue and the α-anomer is purple. A plot over time of the integrals of the peaks corresponding to α and β anomeric protons is shown in Fig. 6d. *(e)* Representation of the mechanism of endorhamnosidase activity. The orange ellipses are the representation of the amino acids involved in the enzymatic reaction following an inversion mechanism *(f)* Representation of the mutarotation after the endorhamnosidase activity (EA). *(g)* Close-up of $^1$H-NMR spectra from *(a)* of the acetyl region at different time points of the polysaccharide sample in the presence of gp20(248-1070). Signals of polysaccharide acetyl ester group (green) and free acetate released (pink) are highlighted. *(h)* Plot over time of the integrals of the peaks corresponding to the disappearance of the acetyl protons (green) from position 6 of Gal and the appearance of the CH3 proton (pink) from the acetic acid shown in *(g)*. The y-axis corresponding integrated area of NMR peaks has arbitrary units. *(i)* Representation of the esterase activity in the ester of galactose. This enzyme hydrolyses the galactose acetyl ester into free acetic acid and free hydroxymethylene exocyclic group of galactose.

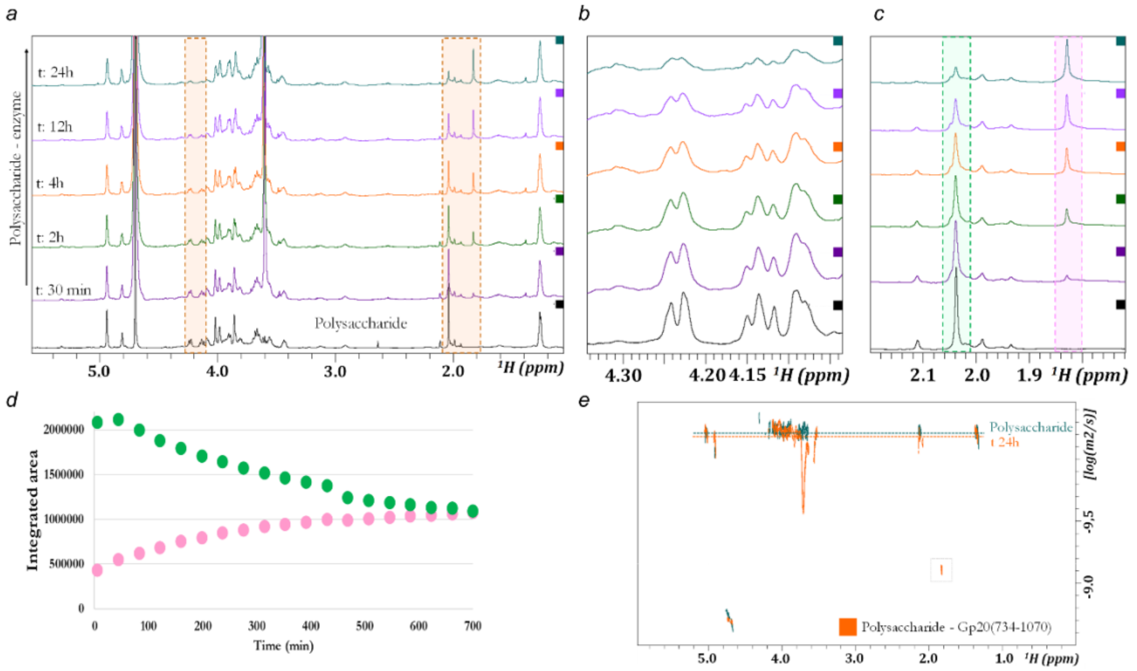


**Figure S6. Enzymatic activity of gp20(734-1070) on the Anatum polysaccharide**

*(a)* $^1$H-NMR spectra of the polysaccharide in the presence of gp20 (734-1070) at different times. The areas with changes are marked in the dashed brown rectangle. The gp20(734-1070) concentration was 1 µM, with a 50:1 ligand protein ratio. *(b-c)* Enlargement of areas from *(a)* that change over time due to esterase activity in the $^1$H-NMR spectra. *(b)* corresponds to the 6-O-acetylated exocyclic $CH_2$ of galactose and *(c)*, to the bound (piacetate ester group. *(d)* Time representation of the disappearance of the acetyl protons (green) from position 6 of Gal and the appearance of the CH3 protons signal (pink) from the acetic acid. The y-axis corresponding to the integrated area of NMR peaks has arbitrary units. *(e)* DOSY spectra of the comparison of polysaccharide size in the presence and absence of the enzyme. The decrease in molecular size is minimal as the polysaccharide chain is not cleaved with the esterase reaction.

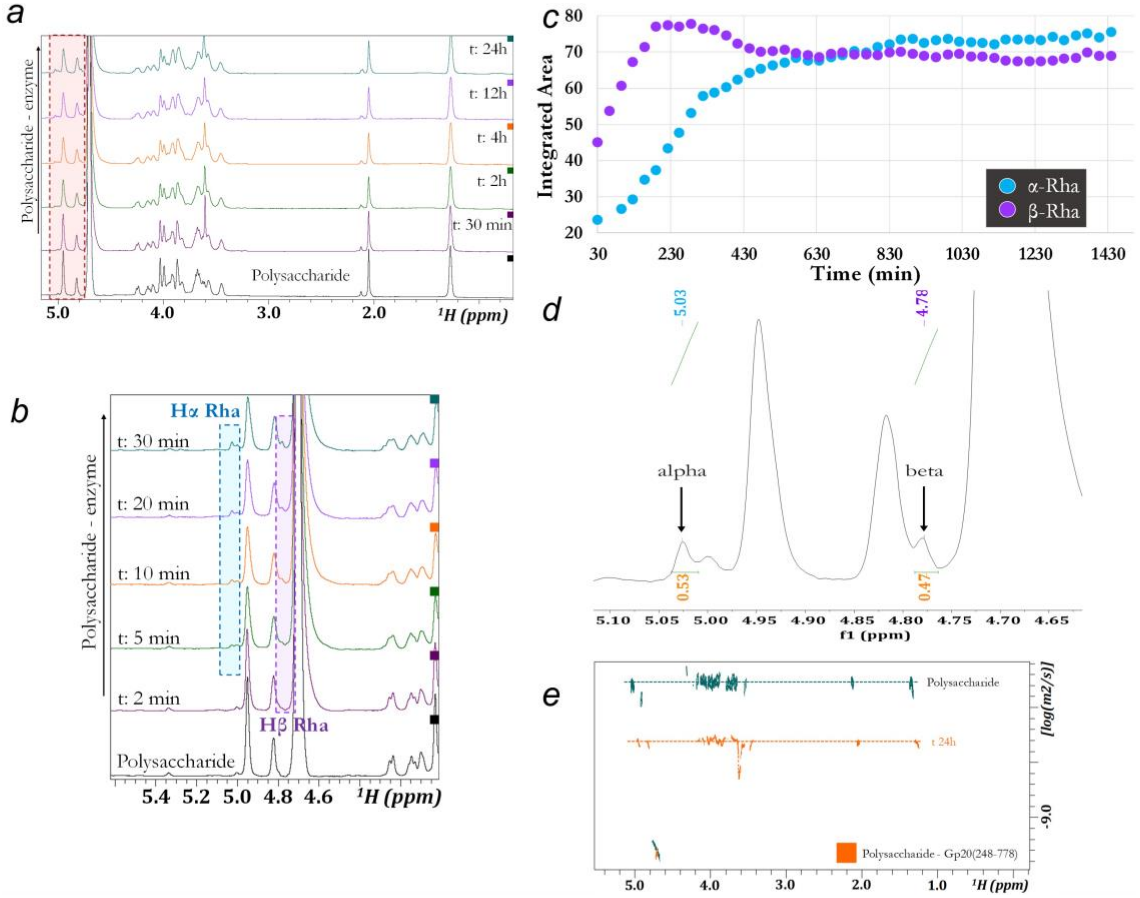


**Figure S7. Enzymatic activity of gp20(248-778) on the Anatum polysaccharide**

*(a)* Areas of change marked in the $^{1}$H-NMR spectra. The gp20(248-778) was at 0.5 µM and a ligand to protein ratio of 50:1. *(b)* Assignment of the two new signals formed in the anomeric region to the anomeric region. The gp20(248-778) was at 0.5 µM and with a 50:1 ligand protein ratio. *(c)* Plot over time (min) of the presence and equilibrium of the alpha and beta isomers of the formed rhamnose. The y-axis corresponding integrated area of NMR peaks has arbitrary units. *(d)* $^{1}$H-NMR spectrum of the polysaccharide in the presence of the enzyme (gp20(248-778)) after 48 hours. The spectrum was analysed with MNova (Mestrelab). *(e)* 2D DOSY spectra of polysaccharide in the presence (orange) and absence (green) of enzyme.

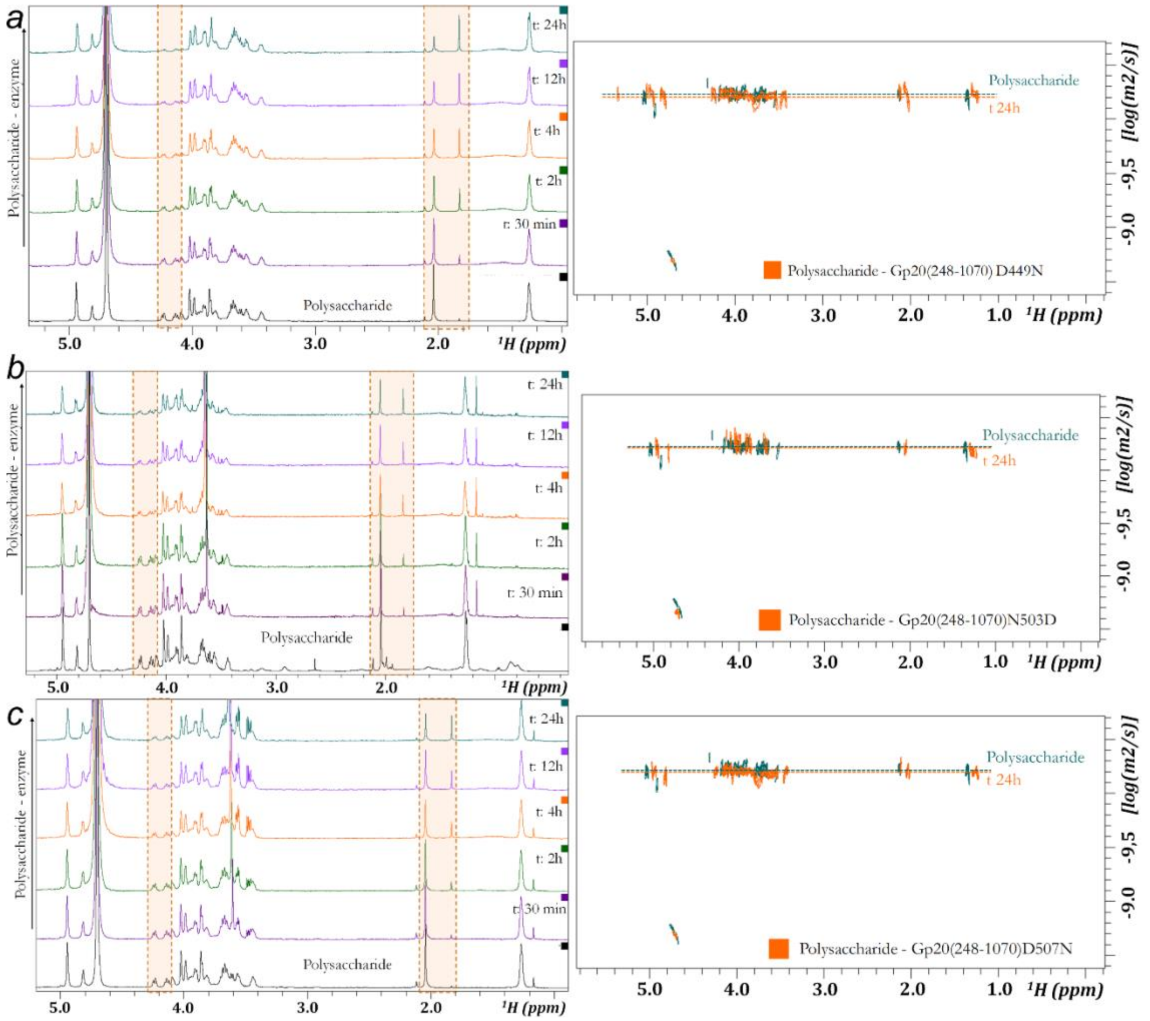


**Figure S8. Enzymatic activity of gp20(248-1070) endorhamnosidase mutants on the Anatum polysaccharide**

*(a-c)* The $^{1}$H-MRN (left) spectra and DOSY (right) of the polysaccharide in the absence/presence of gp20(248-1070) three mutants, D449N *(a)*, N503D *(b)*, D507N *(c)*. The dashed rectangle marks the area of change over time. The enzyme concentration was at 2 μM and a ligand to protein ratio of 50:1.

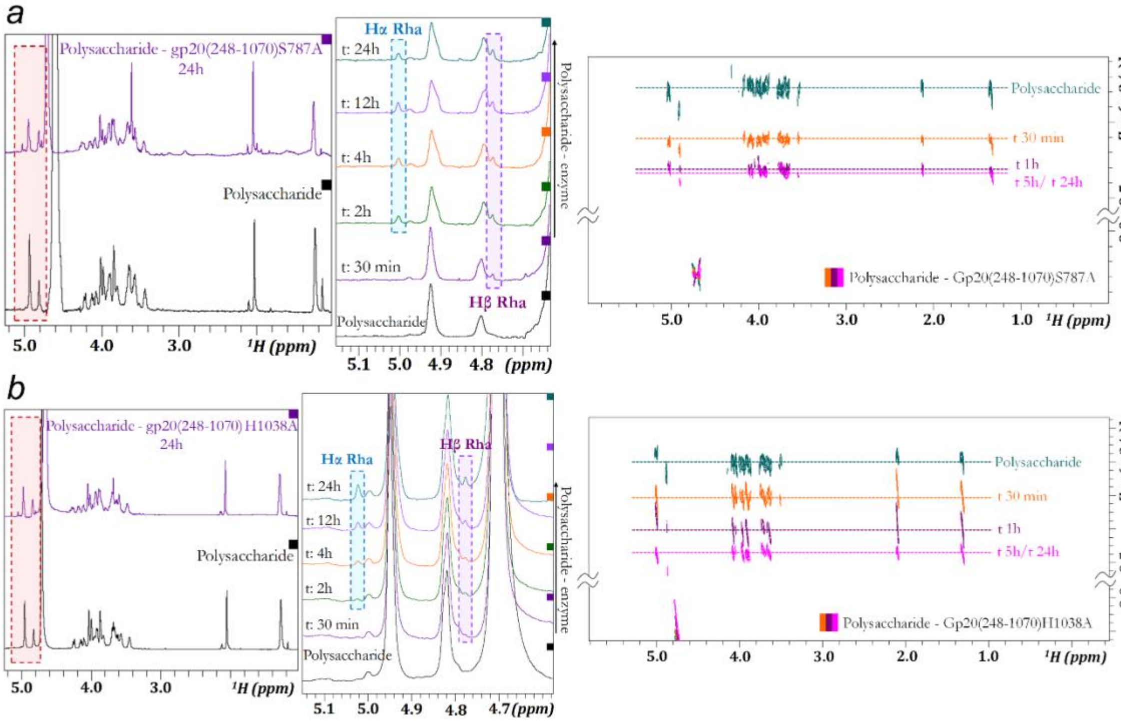


**Figure S9. Enzymatic activity of gp20(248-1070) esterase mutants on the Anatum polysaccharide**

*(a-b)* The $^{1}$H-MRN (left) spectra, its close-up (middle) and DOSY (right) of the polysaccharide in the absence/presence of gp20(248-1070) two mutants, S787A *(a)* and H1038A *(b)*. The dashed rectangle marks the area of change over time. The enzyme concentration was at 2 µM and a ligand to protein ratio of 50:1.

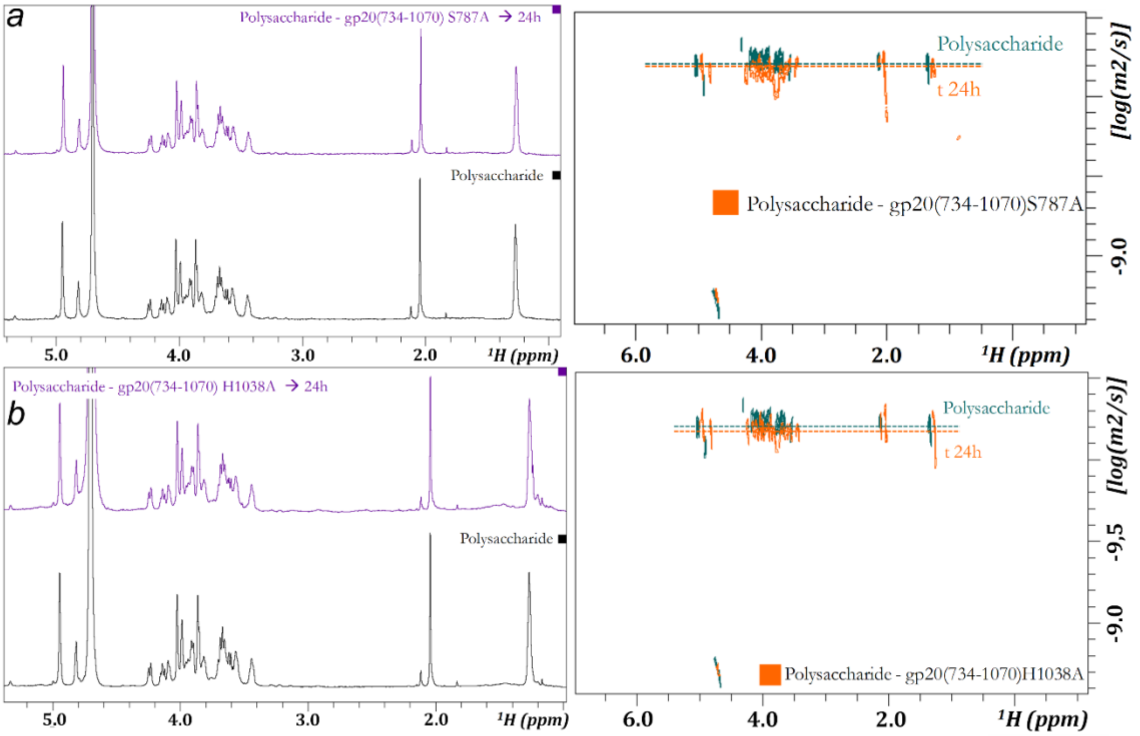


**Figure S10. Enzymatic activity gp20(734-1070) esterase mutants on the Anatum polysaccharide**

The $^{1}$H-NMR (left) and DOSY (right) spectra of the polysaccharide in the presence of two mutants of gp20 (734-1070), S787A *(a)* and H1038A *(b)*, over a 24 h period. The enzyme concentration was at 2 µM and a ligand to protein ratio of 50:1.

| Proton | $^{1}$H Chemical shift | $^{13}$C Chemical shift |
|---|---|---|
| H1 Gal | 4.94 | 98.2 |
| H2 Gal | 3.89 | 70.17 |
| H3 Gal | 3.85 | 77.51 |
| H4 Gal | 3.98 | 69.03 |
| H5 Gal | 4.08 | 68.52 |
| H6 Gal Oac | 4.23 | 63.88 |
| H6´ Gal Oac | 4.13 | |
| CH3 Gal Oac | 2.04 | 20.11 |
| H6 Gal OH | 3.66 | 60.94 |
| H6’ Gal OH | 3.63 | |
| H1 Man | 4.81 | 101.01 |
| H2 Man | 4.02 | 70.68 |
| H3 Man | 3.55 | 73.22 |
| H4 Man | 3.85 | 67.51 |
| H Man | 3.43 | 74.39 |
| H6 Man | 3.94 | 66.07 |
| H6´ Man | 3.92 | 80.09 |
| H1 Rha | 4.94 | 102.24 |
| H2 Rha | 3.98 | 70.29 |
| H3 Rha | 4.02 | 68.88 |
| H4 Rha | 3.59 | 80.09 |
| H5 Rha | 3.81 | 67.82 |
| CH3 Rha | 1.25 | 17.18 |

**Table S1. $^{1}$H-$^{13}$C NMR chemical shift (δ, ppm) of polysaccharide.**
